# Insights into the molecular systematics of montane voles of the Western Himalayan Mountains

**DOI:** 10.64898/2026.10.02.756046

**Authors:** Muhammad Raza Khan, Muhammad Sajid Nadeem, Jeffrey L. Coleman, David C. Cannatella

**Affiliations:** Department of Zoology, Wildlife and Fisheries, PMAS-Arid Agriculture University Rawalpindi, Pakistan; Department of Integrative Biology and Biodiversity Center, University of Texas, Austin, TX 78712, United States

**Author notes:** Corresponding author: Department of Integrative Biology, Patterson Labs, 2415 Speedway, University of Texas, Austin, TX 78712, United States.

**Keywords:** *Alticola*, Arvicolinae, Cytb, Himalaya, India, *Hyperacrius*, Pakistan, voles

## Abstract

Kashmir voles (*Hyperacrius*) are poorly studied fossorial arvicolines of the southern Himalayan slopes, with two species recognized. We sequenced Cytb from voles collected in northern Pakistan and combined these data with existing GenBank sequences, yielding 194 tips spanning nearly all arvicoline genera. We analyzed five data sets with maximum likelihood and compared external measurements among species using principal component and linear discriminant analyses.

We recovered five well-supported clades of *Hyperacrius* separated by ∼12–14% p-distance, which we treat as distinct species. We recognize clade 1 as *H. wynnei*, and populations referred to *H. w. traubi* are genetically indistinguishable from *H. w. wynnei*. We recognize *H. zygomaticus*, new rank (clade 2), and tentatively allocate clade 5 to *H. brachelix*. Clades 3 and 4 likely represent undescribed species. Morphometric analyses of external measurements clearly distinguish *H. wynnei* but not clades 3 and 4 (overall classification accuracy 70.3%).

Phylogenetic analyses including nuclear genes placed *Hyperacrius* as sister to a clade containing Microtini, *Arvicola*, Lagurini, *Dinaromys*, and Ellobiusini, whereas mitochondrial analyses weakly allied it with Ellobiusini. Its traditional placement in Clethrionomyini is thus unsupported, although its exact position remains unresolved. *Alticola* is nonmonophyletic, with *A. macrotis* sequences nested within *Clethrionomys*, likely reflecting mitochondrial capture.

**Teaser Text:** Kashmir voles, burrowing rodents of the Himalayan foothills, were thought to include two species. DNA evidence reveals at least five, including two likely new to science, and shows these voles are not close relatives of the red-backed voles they were long grouped with.

## Introduction

Mountains shape climates, create varied habitats, and act as cradles, barriers, bridges, and reservoirs for species, and therefore are a major contributor to biodiversity (Rahbek et al. 2019; Perrigo et al. 2020). In particular, the biodiversity of alpine regions is poorly characterized because of their inaccessibility and logistic challenges for conducting research.

The northern regions of Pakistan, Afghanistan, and India are home to some of the world’s most spectacular mountain ranges, including portions of the Karakoram, Hindu Kush, and Himalayan systems. They also host a remarkable diversity of rodent species in a mosaic of habitats, from alpine meadows and coniferous forests to steppe grasslands and arid rocky slopes (Roberts 1997). These rodents are seed dispersers, soil aerators, and key prey for carnivores such as foxes, raptors, and mustelids. Alpine rodent communities include a mix of Palearctic, Central Asian, and Himalayan elements, with species adapted to both low-elevation valleys and high-altitude zones above 4,000 meters. Endemism is likely relatively high in these isolated valleys, although this has not been examined. The rodent fauna of these regions remains important for biodiversity research, both as foundational science and for understanding ecosystem health in a region bridging several of Asia’s major biogeographic zones.

Common taxa include voles (*Microtus* and *Alticola*), hamsters (*Cricetulus*), and mice (*Apodemus* and *Mus)*. Voles belong to the subfamily Arvicolinae (Rodentia: Cricetidae), which also includes muskrats and lemmings. Arvicolines (33 genera with 182 species [Mammal Diversity Database 2025]) are a young, rapidly radiating, and highly diverse group of species that inhabit almost all terrestrial habitats in temperate and cold climates throughout the Northern Hemisphere. Molecular phylogenetic analyses, mostly of the Cytb gene, have led to a series of taxonomic revisions (Galewski et al. 2006; Lebedev et al. 2007; Robovský et al. 2008; Abramson et al. 2009; Bannikova et al. 2010; Fabre et al. 2012; Liu et al. 2012; Kohli et al. 2014; Pradhan et al. 2019). Arvicolines are also well represented in the Asian fossil record (Fejfar et al. 2011; Withnell and Scarpetta 2024).

The Kashmir voles, *Hyperacrius*, are particularly poorly studied. These fossorial voles are found in pine forests, adjacent fields, pastures, and alpine meadows (1950–3660 m [6400 to 12,000 ft]). The species occur exclusively along the southern Himalayan slopes in northeastern Pakistan, and India (Phillips 1969; Pradhan and Talmale 2009). At higher elevations, these voles serve as potential hosts of scrub typhus (Traub and Evans 1967; Traub et al. 1967).

Miller (1896) established *Hyperacrius* as a subgenus of *Microtus* with type species *Arvicola fertilis* True 1894, viewing them as fossorial descendants of *Alticola*, a high-elevation group, based on cranial and dental characters reflecting burrowing adaptations. Hinton (1926) elevated *Hyperacrius* to generic rank. Phillips (1969) provided the most comprehensive taxonomic treatment. Two species are recognized, *H. fertilis* and *H. wynnei*. They are generally parapatric except at Shogran, Pakistan, where they occur within meters (Traub and Evans 1967). *Hyperacrius fertilis* has three subspecies: *H. f. fertilis* (True 1894), *H. f. zygomaticus* Phillips 1969, and *H. f. brachelix* (Miller 1899). *Hyperacrius wynnei* has two subspecies: *H. w. wynnei* (Blanford 1881) and *H. w. traubi* Phillips 1969.

Here we infer a phylogeny of newly collected samples of Kashmir voles (*Hyperacrius*) as well as ancillary samples of mountain voles (*Alticola*) and integrate our data with existing DNA sequences from nearly all extant arvicoline genera. We address species diversity of *Hyperacrius* and interrelationships among arvicolines, with reference to *Hyperacrius* and *Alticola*. A detailed critique of arvicoline phylogeny is beyond the scope of this paper.

## Methods

### Study area

Field work was conducted in the northern mountainous regions of Pakistan; specimens were collected by MRK from April 2021 to May 2023. Collecting localities, selected localities from the literature, and coordinates are given in the Appendix, Figure 1, and Supplementary Data SD1. “Yakhtangay Shangla” is occasionally shortened to Yakhtangay.

**Fig. 1.**
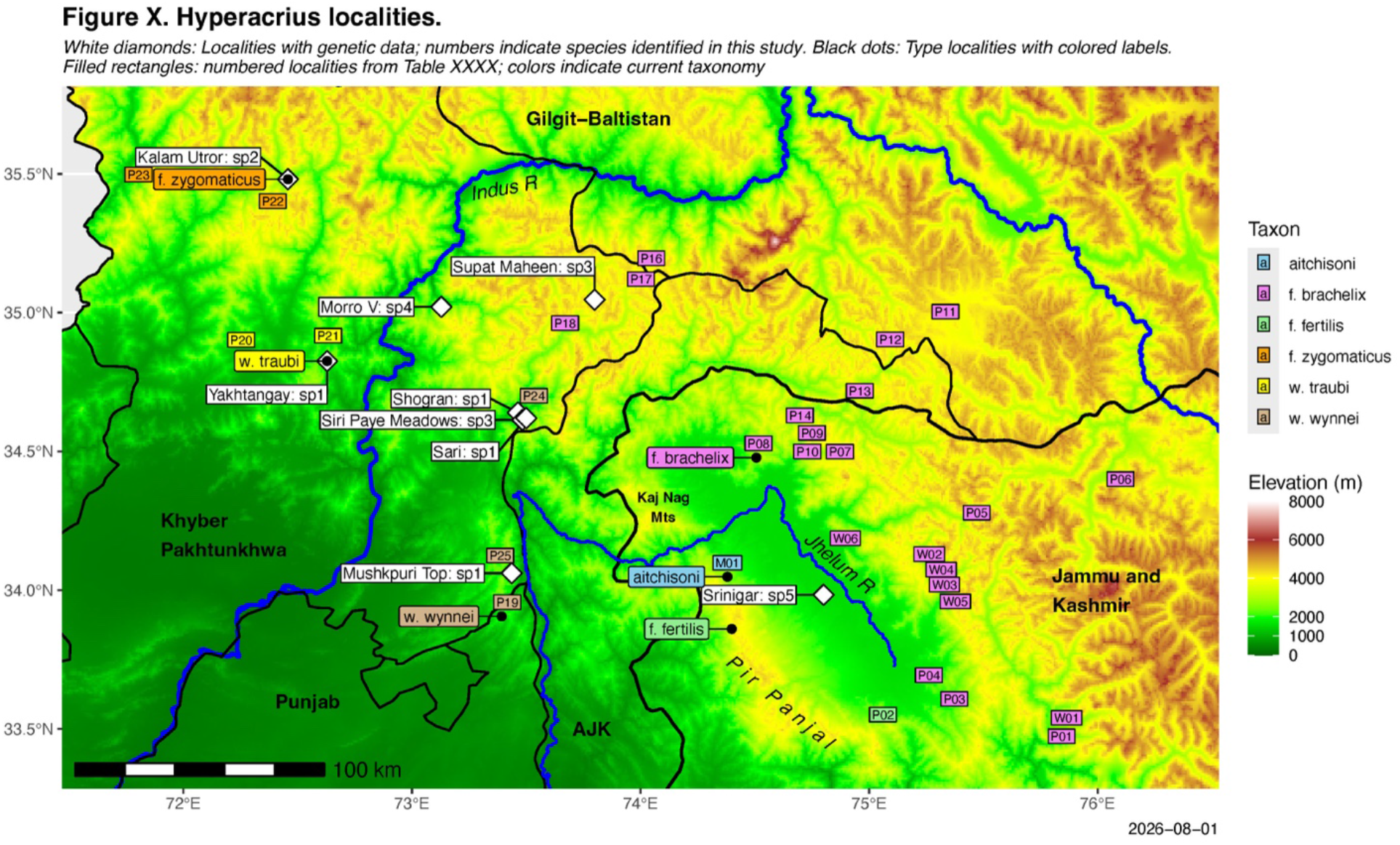
Map of Pakistan and India, showing sampling sites for sequences used in this study in white boxes with numbered species designation as in Fig. 2. Type-localities of nominal taxa are indicated by black dots with labels colored by taxon according to current taxonomy. Numbers for literature localities are shown in small rectangles colored by taxon. For these, “P” indicates numbered localities in Phillips (1969), “W” indicates selected collecting localities of A. E. Ward, and “M” indicates the type-locality of *Hyperacrius aitchisoni*. AJK = Azad Jammu and Kashmir. See details in Supplementary Data SD1.

The voles were captured using mouse traps baited with peanut butter. *Apodemus* (field mice), *Alticola* (mountain voles) and *Mus* were collected incidentally. Guidelines of the American Society of Mammalogists were followed for trapping specimens. Tail tips were removed and preserved in 100% ethanol for DNA extraction. Specimens are stored in the Collections of the Department of Zoology Wildlife and Fisheries, PMAS-AAUR, and all animal procedures were done following institutional approval at the Department of Zoology Wildlife and Fisheries, PMAS-AAUR.

### Molecular methods

DNA was extracted from the tail samples with QIAGEN DNeasy Blood & Tissue kits according to the manufacturer’s protocol. The mitochondrial Cytb gene (Cytb) was amplified using PCR. We prepared a total volume of 50 µl PCR reaction, comprising 10 µl of Q5 Buffer, 1 µl of mM dNTPs, 2.5 µl of forward primer = GACATGAAAAATCATCGTTGTTATT (L14728 of Lebedev et al. 2007) and 2.5µl of reverse primer = ACTGGTTTACAAGACCAGTGTAAT (H15906arvic of Lebedev et al. 2007), 0.5 µl of Q5 hot-start DNA polymerase, and 50–100 ng/µl of freshly extracted DNA template, the rest of the volume was filled with DNA/RNA-free water for a total of 50µl. PCR products were separated in a 1% agarose gel and purified and Sanger-sequenced by Eton Bioscience, Inc. Both strands were sequenced.

### Phylogenetic analysis

We generally follow the Mammal Diversity Database (2025) for the generic taxonomy and Withnell and Scarpetta (2024) for taxonomy at the tribe level (Supplementary Data SD2), except for the recognition of *Aschizomys* as a genus rather than a subgenus of *Alticola*, and recognition of *Alexandromys* and *Stenocranius* as genera rather than subgenera of *Microtus*.

Geneious Prime 2024.1 (Biomatters 2024) was used to edit sequences and Mesquite 4 (Maddison and Maddison 2026) and MAFFT v7.475 (Katoh and Standley 2013) were used for alignment and determination of codon positions. We detected no evidence of numts (Triant and DeWoody 2008), based on codon structure.

We used the “combined.nex” matrix of Withnell and Scarpetta (2024), which included five genes (Cytb, COI, Ghr, IRBP/RBP3, and BRCA1; 5847 sites) and 149 sequences. All sequences were analyzed with BLAST (Camacho et al. 2009) to confirm species identity and a few problematic sequences were removed.

We augmented this with Cytb sequences from GenBank and our new Cytb sequences of *Hyperacrius* and ancillary samples of *Alticola.* This matrix had 5857 sites and 194 tips (Supplementary Data SD3). Five ML analyses were performed: Analysis 1 with all five genes, Analysis 2 with mtDNA only, Analysis 3 with Cytb only, Analysis 4 with COI only, and Analysis 5 with Cytb and three nuclear genes. Some corrections were made to the matrix (Supplementary Data SD4). The modified list of GenBank sequences from Withnell and Scarpetta (2024) is Supplementary Data SD5.

All maximum-likelihood analyses were performed using IQ-TREE 3 v3.1.1 (Wong et al. 2026). ModelFinder (Kalyaanamoorthy et al. 2017), implemented in IQ-TREE, was used to find the best model of molecular evolution for each data partition under the Bayesian Information Criterion (BIC) with subsequent merging (option: -m TESTMERGE), restricting rate heterogeneity to a gamma distribution and excluding models with invariable sites (-m I). Base frequencies were estimated by maximum likelihood (+FO). The models were edge-linked (-p) (Chernomor et al. 2016). In all analyses, protein-coding genes were partitioned by codon position using Mesquite 4 (Maddison and Maddison 2026).

Branch support was assessed using ultrafast bootstrapping (Minh et al. 2013). We ran five independent searches, each with 50,000 ultrafast bootstrap (Hoang et al. 2018) replicates (options: --runs 5 -B 50000) with a broad branch-swapping radius (--radius 180). FigTree 1.4.1 (Rambaut 2007) was used to visualize trees. We used R scripts with dist.dna() from the ape package (Paradis and Schliep 2019) to calculate the uncorrected p-distance (proportion of sites at which two sequences differ) to quantify genetic divergence.

### Morphometrics

Standard measurements for rodents (Hall 1981) were taken (Supplementary Data SD6). These were head–body length (HBL), tail length (TAIL), hindfoot length (HFL), and ear length (EL). Total Length (TL) was calculated as the sum of HBL and TAIL. Therefore, TL is not independent of those two variables, and so we used HBL as an estimate of size.

Most statistical analyses were done in R (R Core Team 2025) using RStudio (Posit Team 2026) and the base package. Boxplots for mensural characters and statistical tests were produced using R code (Supplementary Data SD7). A Kruskal–Wallis test (kruskal.test) was performed to assess whether the median value of at least one of the four species was significantly different from the others, followed by a Dunn test (Dinno 2024) of all pairwise comparisons with Benjamini–Hochberg adjustment for false positives (dunn.test). Based on the Dunn test, letters were used on the plot to indicate which species were significantly different for a variable; boxplots that share a letter are not significantly different at a type I error rate of α = 0.05. Principal component analysis (PCA) was done with prcomp() on raw variables with a correlation matrix. The linear discriminant analysis (LDA) was done using MASS::lda() with species as a priori groups; a cross validation analysis was done using “leave-one-out” with equal priors.

## Results

### Species delimitation in *Hyperacrius*

Our samples represent the largest phylogenetic dataset for this genus to date. All phylogenetic analyses (Fig. 2) indicate the presence of five well-supported clades of *Hyperacrius*, which we treated as full species based on the level of genetic divergence (see Discussion). When possible, we allocate available names to the five clades and under the Discussion we justify our taxonomic decisions, using both geographic distribution and proximity of type-localities, phenotype, and phylogenetic relationships.

**Fig. 2.**
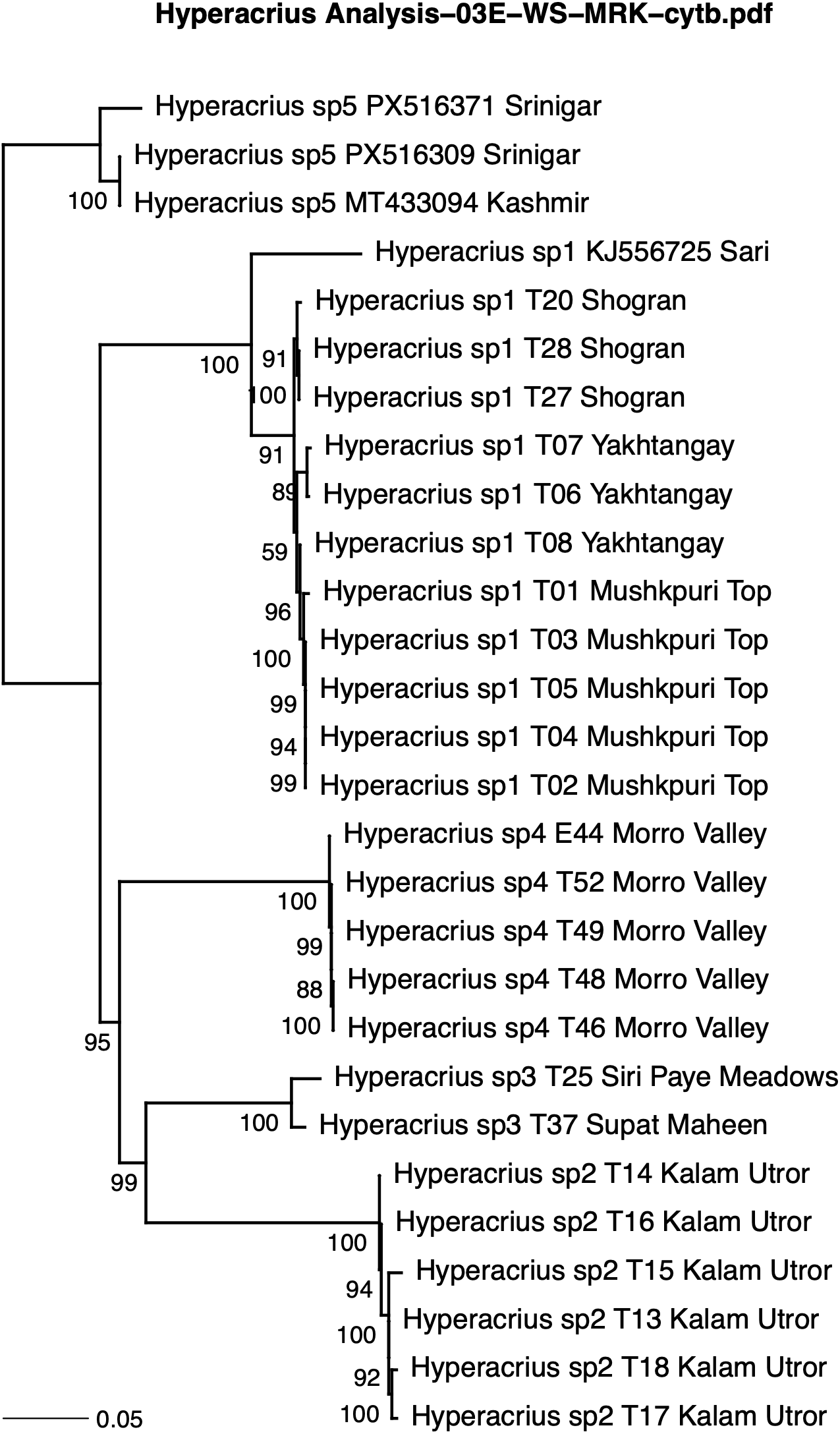
Maximum likelihood phylogeny of *Hyperacrius* taken from Analysis 3 (Cytb). The scale units are substitutions per site. Ultrafast bootstrap values are shown.

#### Hyperacrius sp. 1

Our samples of *Hyperacrius* sp. 1 are from Shogran (2350 m) in the Kaghan Valley, Mushkpuri Top (2745 m) north of Murree, and Yakhtangay (Figs. 1, 2). The p-distance between most sequences of *Hyperacrius* sp. 1 is <1.4%. The mean distance from the GenBank sequence KJ556725 (Kohli et al. 2014) from Sari, Pakistan, to the other samples of *H.* sp. 1 is ∼6.0%, but examination of the alignments suggests sequence quality problems, so we regard this large distance as an artifact, and we consider this sequence to belong to *H*. sp. 1.

The mean pairwise p-distance from *H.* sp. 1 to other species of *Hyperacrius* is 11.9%–13.4% (Table 1), which suggests that this taxon should be treated as a species rather than subspecies (see Discussion).

**Table 1.** Uncorrected p-distances and K2P distances among species of *Hyperacrius*.

| Species | Species | Mean | Minimum | Maximum | Median |
| --- | --- | --- | --- | --- | --- |
| Uncorrected p-distance |  |  |  |  |  |
| sp. 1 | sp. 1 | 0.0068 | 0.000 | 0.014 | 0.008 |
| sp. 1 | sp. 2 | 0.1340 | 0.130 | 0.140 | 0.134 |
| sp. 1 | sp. 3 | 0.1210 | 0.118 | 0.125 | 0.121 |
| sp. 1 | sp. 4 | 0.1190 | 0.112 | 0.126 | 0.119 |
| sp. 1 | sp. 5 | 0.1210 | 0.116 | 0.123 | 0.122 |
| sp. 2 | sp. 2 | 0.0064 | 0.000 | 0.012 | 0.007 |
| sp. 2 | sp. 3 | 0.1300 | 0.128 | 0.133 | 0.129 |
| sp. 2 | sp. 4 | 0.1380 | 0.133 | 0.147 | 0.137 |
| sp. 2 | sp. 5 | 0.1340 | 0.131 | 0.137 | 0.135 |
| sp. 3 | sp. 3 | 0.0118 | 0.000 | 0.024 | 0.012 |
| sp. 3 | sp. 4 | 0.1280 | 0.122 | 0.134 | 0.127 |
| sp. 3 | sp. 5 | 0.1240 | 0.120 | 0.127 | 0.125 |
| sp. 4 | sp. 4 | 0.0003 | 0.000 | 0.001 | 0.000 |
| sp. 4 | sp. 5 | 0.1360 | 0.128 | 0.141 | 0.137 |
| sp. 5 | sp. 5 | 0.0119 | 0.000 | 0.024 | 0.012 |
| K2P distance |  |  |  |  |  |
| sp. 1 | sp. 1 | 0.0069 | 0.000 | 0.014 | 0.008 |
| sp. 1 | sp. 2 | 0.1520 | 0.146 | 0.161 | 0.152 |
| sp. 1 | sp. 3 | 0.1360 | 0.132 | 0.140 | 0.136 |
| sp. 1 | sp. 4 | 0.1330 | 0.124 | 0.141 | 0.133 |
| sp. 1 | sp. 5 | 0.1360 | 0.129 | 0.138 | 0.137 |
| sp. 2 | sp. 2 | 0.0064 | 0.000 | 0.013 | 0.007 |
| sp. 2 | sp. 3 | 0.1470 | 0.144 | 0.151 | 0.145 |
| sp. 2 | sp. 4 | 0.1570 | 0.150 | 0.167 | 0.156 |
| sp. 2 | sp. 5 | 0.1520 | 0.148 | 0.156 | 0.153 |
| sp. 3 | sp. 3 | 0.0121 | 0.000 | 0.024 | 0.012 |
| sp. 3 | sp. 4 | 0.1440 | 0.137 | 0.152 | 0.143 |
| sp. 3 | sp. 5 | 0.1400 | 0.134 | 0.143 | 0.141 |
| sp. 4 | sp. 4 | 0.0002 | 0.000 | 0.001 | 0.000 |
| sp. 4 | sp. 5 | 0.1550 | 0.145 | 0.162 | 0.156 |
| sp. 5 | sp. 5 | 0.0122 | 0.000 | 0.024 | 0.012 |

#### Hyperacrius sp. 2

The largest p-distance among individuals of *Hyperacrius* sp. 2 is 1.2% (mean 0.64%) (Table 1). All samples of *H.* sp. 2 are from Kalam Utror (2407 m), which is a few km from Utror, the type-locality of *Hyperacrius fertilis zygomaticus* Phillips 1969 (Table 2). Utror is a valley in the Upper Swat District in Khyber Pakhtunkhwa province (Fig. 1).

**Table 2.** Type localities and geographic coordinates of nominal taxa of *Hyperacrius*.

| Taxon | Author and Date | Verbatim type-locality | Coordinates |
| --- | --- | --- | --- |
| *Hyperacrius fertilis fertilis* | True 1894 | "Pir Panjal range [...] 8,500 feet [2,600 m]". | 74.3994, 33.8603<br>(estimated for plotting) |
| *Hyperacrius fertilis brachelix* | Miller 1899 | "Nagmarg, Kashmir (altitude, 9,000 feet [2,745 m])" | 74.5068, 34.4776 |
| *Hyperacrius fertilis zygomaticus* | Phillips 1969 | "6 mi SW Utror, 8900 ft [2,710 m], Swat, West Pakistan." | 72.4576, 35.4807<br>(our collection) |
| *Hyperacrius wynnei wynnei* | Blanford 1881 | "Murree, 2,135 m [7000 ft], North Punjab (border of Rawalpindi-Hazara districts) West Pakistan" (as modified by Phillips 1969:462). | 73.3927, 33.9057 |
| *Hyperacrius wynnei traubi* | Phillips 1969 | "Yakh Tangai, 6600 ft [2,010m], Swat, West Pakistan" | 72.6290, 34.8256 |
| *Hyperacrius aitchisoni* | Miller 1897 | "Gulmerg, Kashmir altitude 9000 feet" | 74.3805, 34.0484 |

#### Hyperacrius sp. 3

Our samples of *Hyperacrius* sp. 3 are from Supat Maheen (3618 m) and Siri Paye Meadows (3045 m). The p-distance between these populations is 2.4%. These localities are at higher elevations than Shogran (2350 m) and Mushkpuri Top (2745 m), where we found *H*. sp. 1. As discussed below, *Hyperacrius* sp. 3 may not be referable to any nominal species.

#### Hyperacrius sp. 4

*Hyperacrius* sp. 4 is from the Morro Valley (2965 m; Fig. 1, Appendix). Museum specimens from this region are rare. This species is highly divergent (11.9–13.8% mean p-distance) from other species (Table 1). It may not be referable to any nominal species.

#### Hyperacrius sp. 5

We have no specimens associated with this species. GenBank sequences (PX516309 and PX516371) of *Hyperacrius* from Srinagar, Rangreth, in the Kashmir Valley (33.9832 N 74.8008 E, 1610 m, fide GenBank records), and a single sequence of *Hyperacrius* (MT433094) from “Kashmir” clustered with 100% ufbs (ultra-fast bootstrap) support. The mean p-distance from this species to the other four species is 12.1–13.6% and the distances between MT433094 and the Srinagar sequences are <3%.

### Morphometrics

#### Interspecific differences

We found that some, but not all, traits vary among the four species. For the species-level comparisons, Kruskal–Wallis tests were significant for TL, HBL, HFL, EL, EL/HBL, and TAIL/HBL, and not significant for TAIL and HFL/HBL; none fell in the marginal range (0.03 < *P* < 0.08) (Fig. 3, Supplementary Data SD9).

**Fig. 3.**
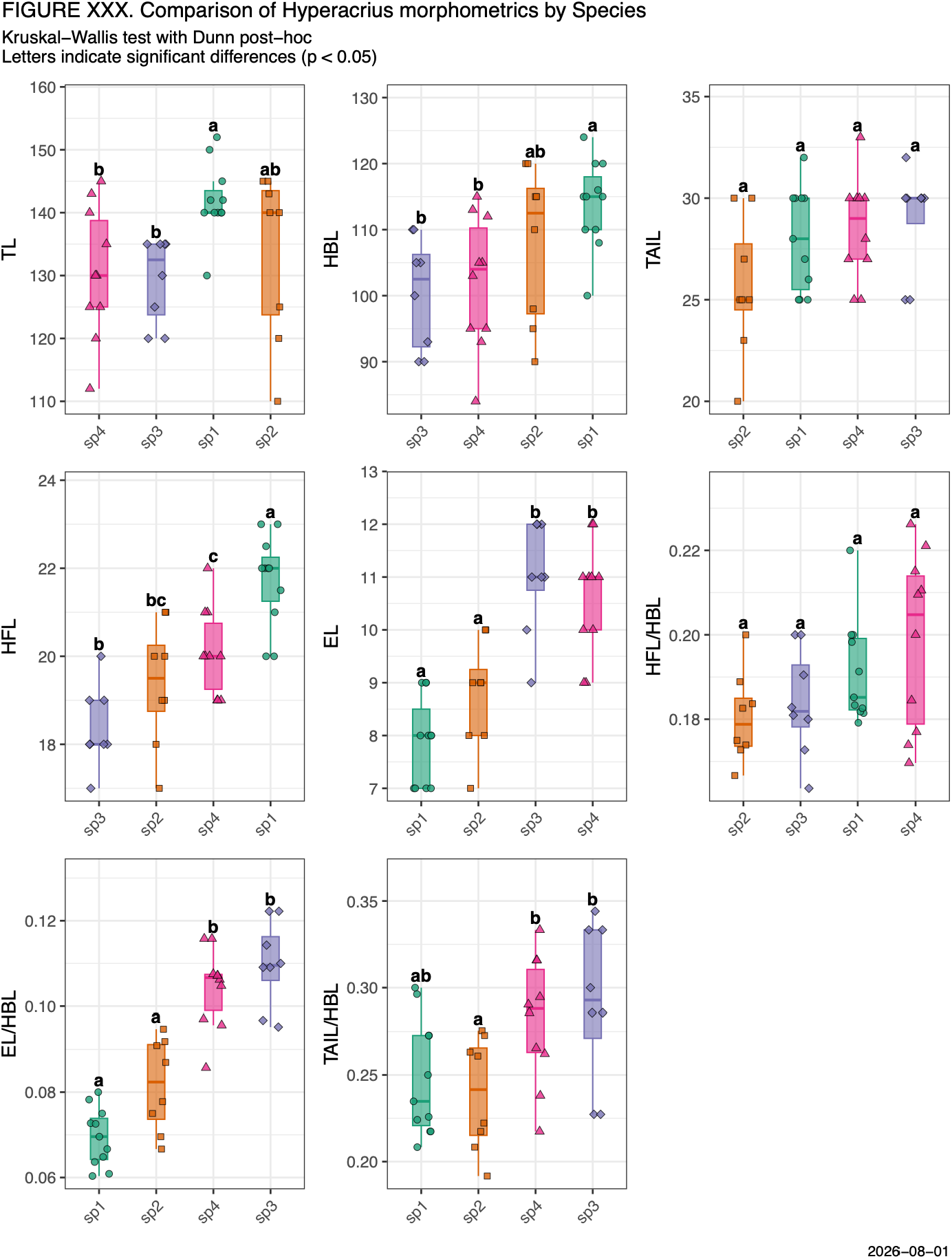
Boxplots of external measurements and ratios by species. For each variable, the Kruskal–Wallis test was used to detect differences among species, and Dunn posthoc tests with Benjamini–Hochberg adjustment were applied to control the false discovery rate. Species that share at least one letter do not differ significantly (*P* ≥ 0.05) for that measurement, and species with no letter in common differ significantly (*P* < 0.05).

Dunn posthoc tests showed that for TL and HBL, *H*. sp. 1 differed from *H*. sp. 3 and *H*. sp. 4. For HFL, *H*. sp. 1 differed from all other species, and *H*. sp. 4 differed from *H*. sp. 3. For EL and for EL/HBL, *H*. sp. 1 and *H*. sp. 2 each differed from *H*. sp. 3 and *H*. sp. 4. For TAIL/HBL, *H*. sp. 2 differed from *H*. sp. 3 and *H*. sp. 4 (Supplementary Data SD9).

#### Principal component analysis

Principal component analysis of the four external measurements (Fig. 4, Table 3, Supplementary Data SD10) did not recover the isometric size axis typical of morphometric datasets. Rather than loading with uniform sign on the first component, the variables partitioned into opposing contrasts: PC1 (explaining 49% of the variance) contrasted HBL and HFL against EL and TAIL, while PC2 (26% of the variance) was dominated by TAIL (correlation loading 0.760). The mixture of positive and negative pairwise correlations among traits indicates that the PC1 from the pooled sample reflects shape rather than overall body size (Supplementary Data SD10).

**Fig. 4.**
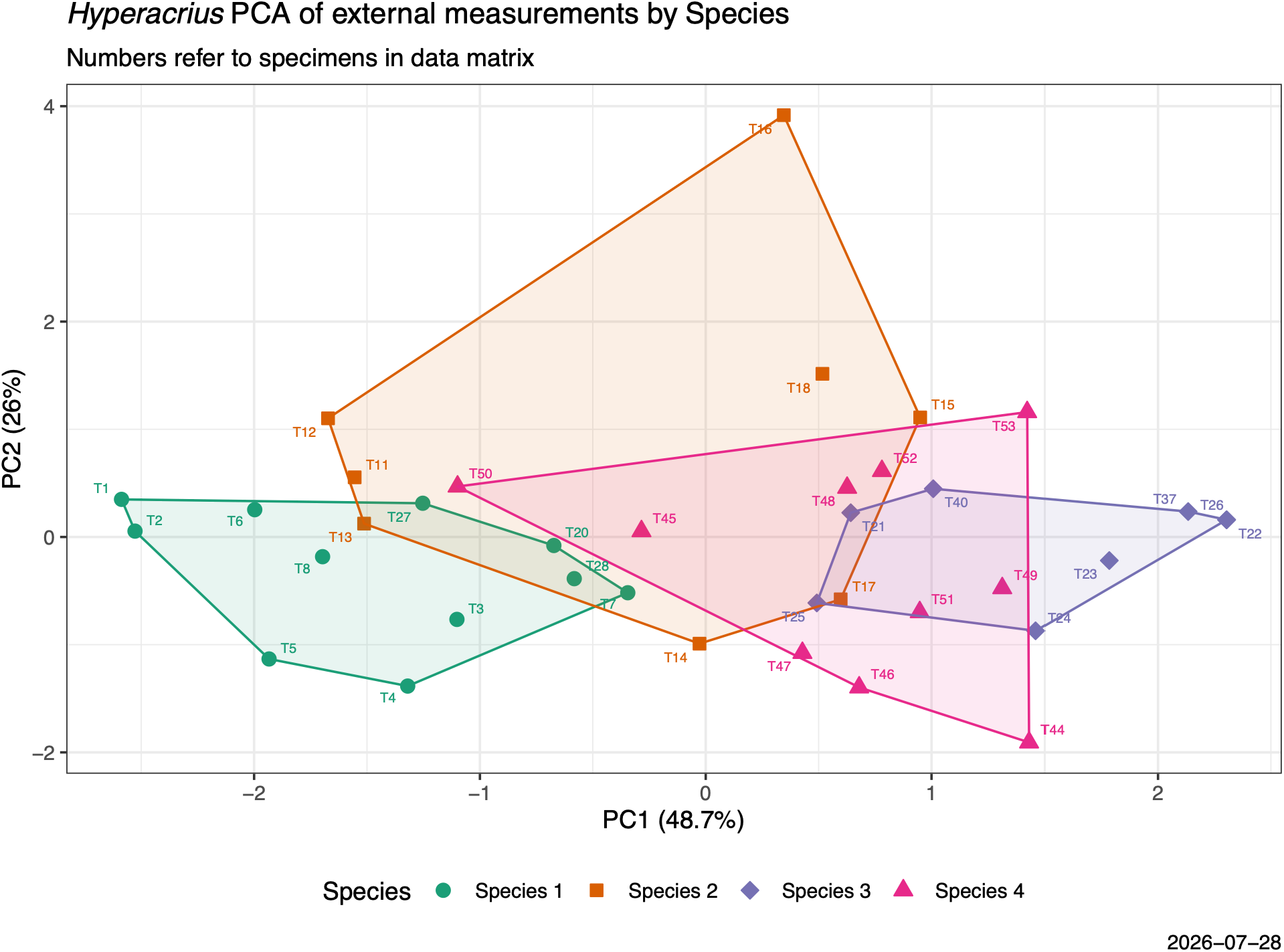
Principal components plot with species circumscribed by convex polygons. Numbered symbols refer to specimens in the data file (Supplementary Data SD6). Axis titles indicate the percent of variance explained by that axis.

**Table 3.** Correlation loadings of variables on the four axes, and the proportion of variance explained by each. Correlation loadings are the simple correlation between each original variable, across specimens, and the scores on a principal component.

| Variable | PC1 | PC2 | PC3 | PC4 |
| --- | --- | --- | --- | --- |
| HBL | -0.760 | -0.404 | -0.403 | -0.311 |
| TAIL | 0.452 | -0.760 | 0.447 | -0.132 |
| HFL | -0.834 | -0.371 | 0.139 | 0.383 |
| EL | 0.684 | -0.399 | -0.573 | 0.209 |
| Proportion of Variance | 0.487 | 0.260 | 0.178 | 0.076 |

The first PC axis clearly separates individuals of *H.* sp. 1 from *H*. sp. 3. There is slight overlap between *H.* sp. 1 and *H*. sp. 4. This is consistent with the Dunn tests, in which *H*. sp. 1 was significantly different from *H*. sp. 3 and *H*. sp. 4 in HBL and HFL.

The second PC axis was uniformly negative across all four measurements, indicative of a general-size axis with TAIL as the main contributor; the remaining traits loaded moderately but consistently (Table 3). This is consistent with the Kruskal–Wallis analysis, which indicated that only TAIL among the four variables lacked a signal of shape contrast; the species were not significantly different in this trait.

#### Linear discriminant analysis

The first two discriminant functions accounted for nearly all of the separation among the four species (Fig. 5, Table 4, Supplementary Data SD11): LD1 explained 84.1% of the between-group variance and LD2 an additional 14.3%, together accounting for about 98%.

**Fig. 5.**
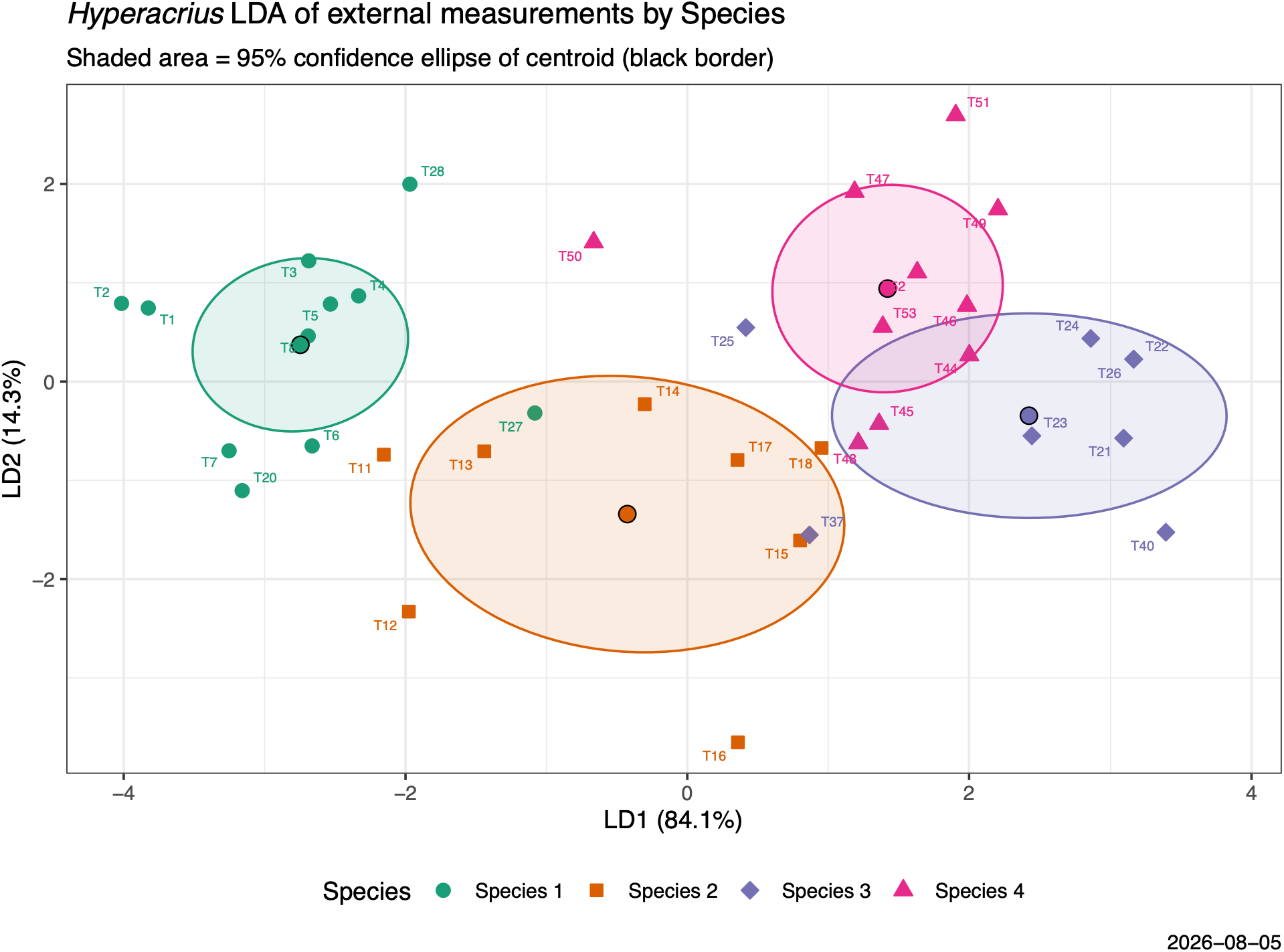
Linear Discriminant Analysis (LDA) plot with species circumscribed by 95% confidence ellipses for centroids. Axis titles indicate the proportion of trace (between-group discrimination) explained by that axis. Centroids are represented by dots with a black border. Numbered symbols refer to specimens in the data file (Supplementary Data SD6)

**Table 4.**
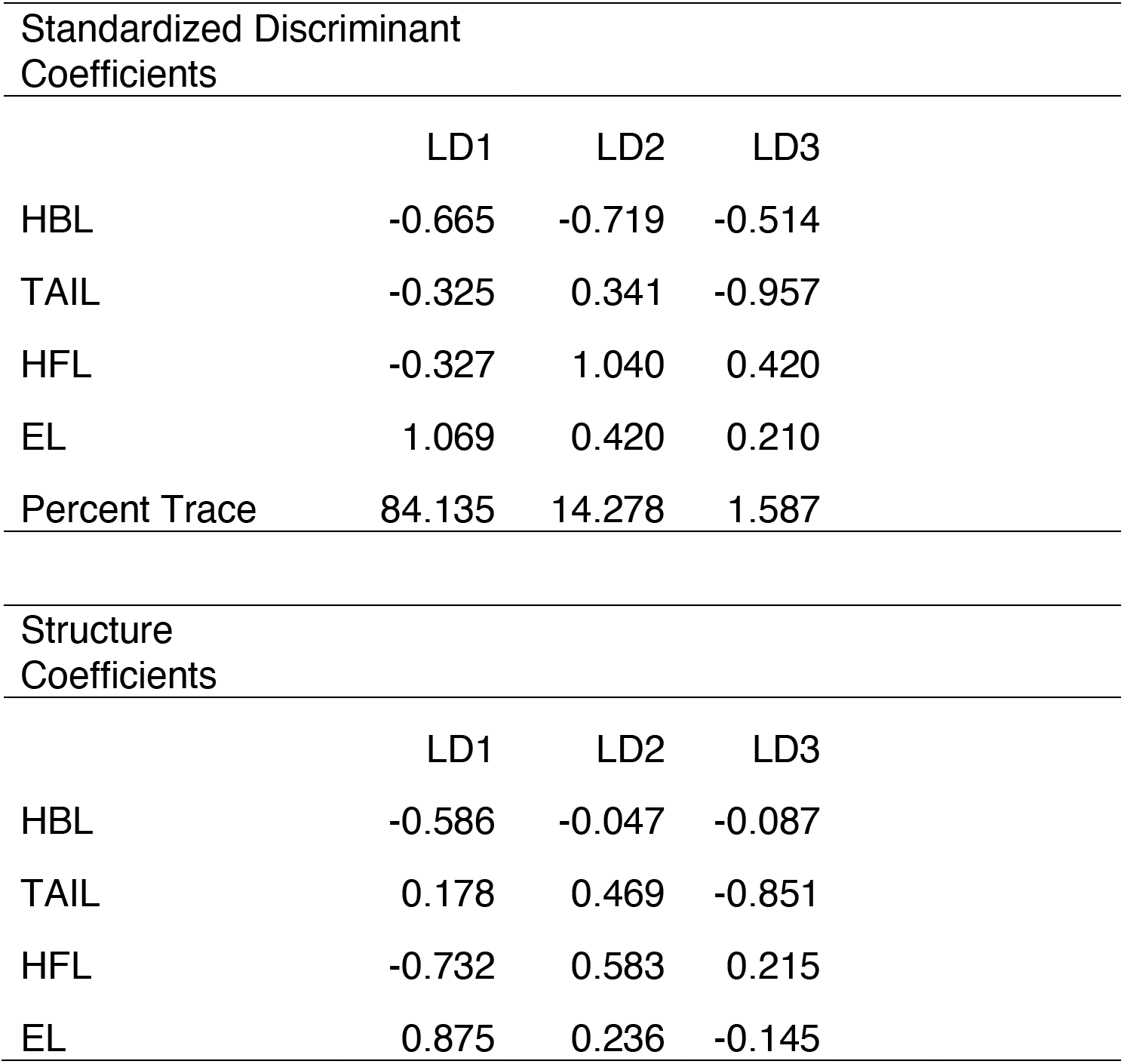
Standardized discriminant coefficients, percent trace, and structure coefficients from the linear discriminant analysis. For the standardized discriminant coefficients, the variable with the largest absolute value contributes most to that axis. Structure coefficients are the simple correlations between each variable and the discriminant scores. Trace is the fraction of total between-group discrimination captured by each discriminant function.

LD1 was defined primarily by ear length (EL), which had by far the largest standardized coefficient (1.07) and structure coefficient (0.88) (Table 4). Head-body length (HBL) and hindfoot length (HFL) loaded negatively (structure coefficients -0.59 and -0.73, respectively). LD2 was defined primarily by HFL (standardized coefficient 1.04, structure coefficient 0.58), with a secondary positive contribution from TAIL (structure coefficient 0.47). HBL also carried a large standardized coefficient on LD2 (-0.72), but its structure coefficient was near zero (-0.05), indicating that HBL is correlated with HFL and TAIL (longer-bodied voles tend to have longer feet and tails too), but that shared variance does not separate species along LD2.

In ordination space (Fig. 5), individuals of *H*. sp. 1 formed a distinct, well-separated cluster at low LD1 values. *Hyperacrius* sp. 2 had a broader, more diffuse distribution overlapping both *H*. sp. 1 and, to a lesser extent, *H*. sp. 3 and *H*. sp. 4. *Hyperacrius* sp. 3 and *H.* sp. 4 both scored high on LD1 and were separated from each other mainly by LD2, but their 95% confidence ellipses overlapped.

The “leave-one-out” cross-validation analysis gave an overall accuracy of 70.3%, well above the 25% expected by chance for four groups (Table 5). Per-species accuracy was uneven. Misclassifications were concentrated between *H*. sp. 3 and *H*. sp. 4 (4 of 10 *H*. sp. 4 individuals were assigned to *H*. sp. 3), while misclassifications of *H*. sp. 2 were split between *H*. sp. 1 (2 individuals) and *H*. sp. 4 (1 individual).

**Table 5.**
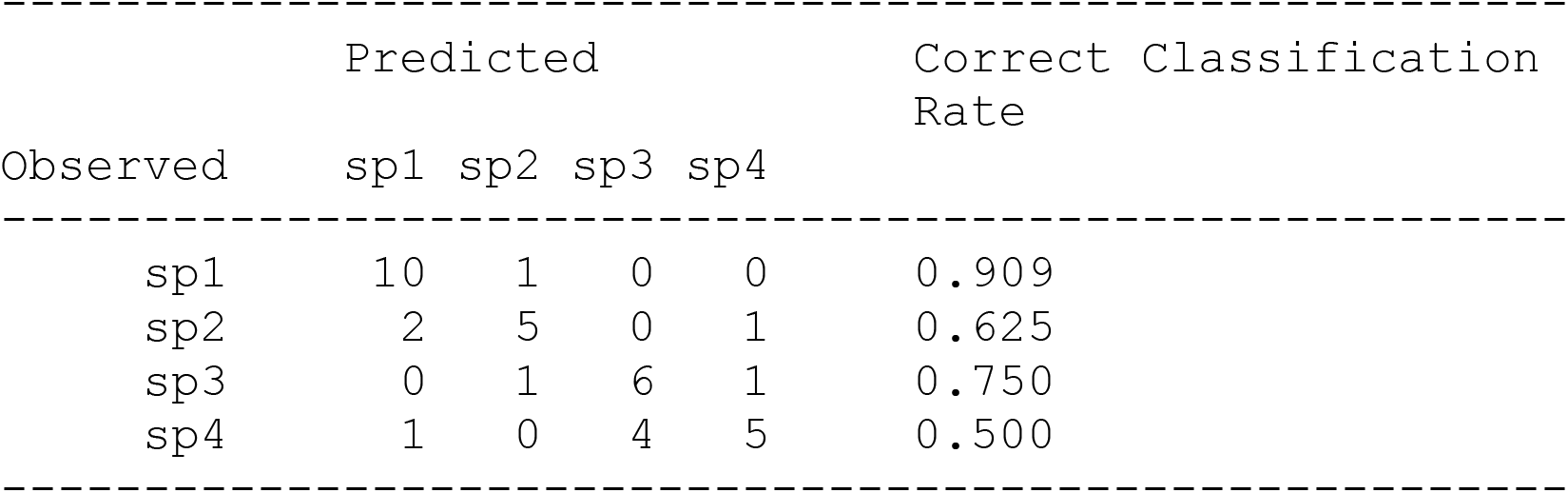
Leave-one-out (LOO) cross-validated classification of *Hyperacrius* specimens from the linear discriminant analysis. Each specimen was classified by a discriminant function computed with that specimen excluded. Rows give observed species assignment and columns give predicted assignment; diagonal entries are the correct classifications. The correct classification rate column is the proportion of specimens of each species assigned to the correct species. Overall cross-validated accuracy was 70.3% (26 of 37 specimens).

### Phylogenetic relationships of *Hyperacrius* to other Arvicolinae

All phylogenetic analyses converged on stable likelihoods in five replicate runs. The summary statistics for the five ML analyses are given in Supplementary Data SD8. In the four analyses that include *Hyperacrius* the relationships of *Hyperacrius* species to each other are identical, with very similar branch lengths and support values. However, the relationships of *Hyperacrius* to other taxa show different degrees of support.

Analysis 1. All five genes (Fig. 6, Supplementary Data SD12). *Hyperacrius* is the sister-group of a poorly supported clade (ufbs = 38) including the Microtini (excluding *Lemmiscus*), *Arvicola*, *Lemmiscus*, Lagurini (*Lagurus* and *Eolagurus*), *Dinaromys*, and Ellobiusini (*Ellobius* and *Bramus*). *Hyperacrius* + this clade is strongly supported (ufbs = 95), and it is the sister group of Clethrionomyini (ufbs = 98).

**Fig. 6.**
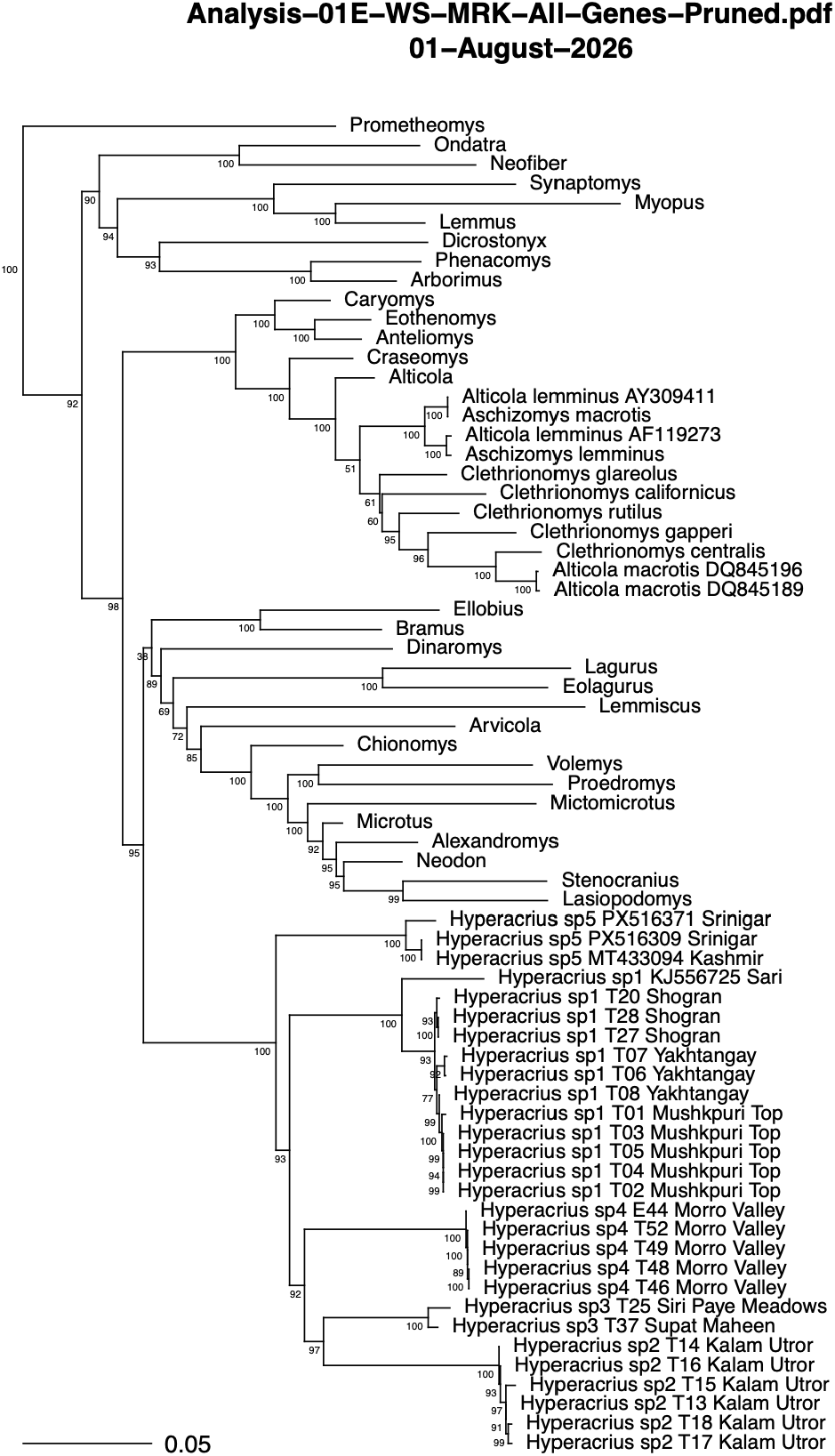
Maximum likelihood phylogeny using mitochondrial and nuclear genes (Analysis 1). Genera with multiple species represented are pruned to the genus name. Ultrafast bootstrap values are shown at the nodes. The scale units are substitutions per site. The complete phylogeny with 194 tips is in Supplementary Data SD12

Analysis 2. mtDNA only (Fig. 7, Supplementary Data SD13). *Hyperacrius* is the sister-group (ufbs = 33) of the Ellobiusini. The closest relatives of this clade are two poorly supported clades including Microtini (excluding *Lemmiscus*), *Arvicola*, *Lemmiscus*, Lagurini (*Lagurus* and *Eolagurus*), and *Dinaromys*. The Clethrionomyini is the sister-group of this larger clade, but ufbs is only 45.

**Fig. 7.**
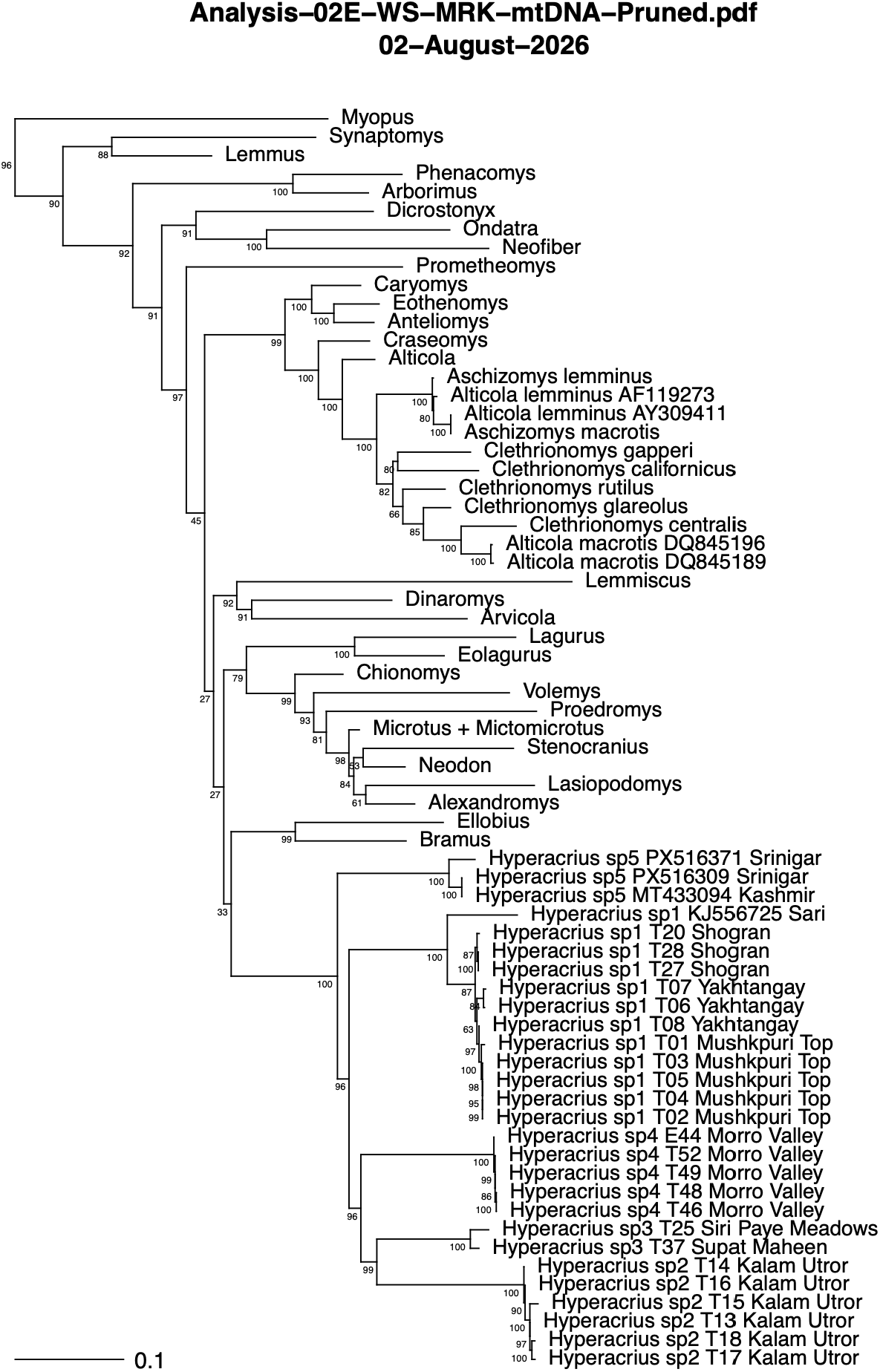
Maximum likelihood phylogeny using mtDNA genes (Analysis 2). Genera with multiple species represented are pruned to the genus name. Ultrafast bootstrap values are shown at the nodes. The scale units are substitutions per site. The complete phylogeny with 194 tips is in Supplementary Data SD13.

Analysis 3. Cytb only (Fig. 8, Supplementary Data SD14). This topology is mostly identical to that of Analysis 2 (mtDNA only), but with slightly higher, but still poor, bootstrap support at most nodes.

**Fig. 8.**
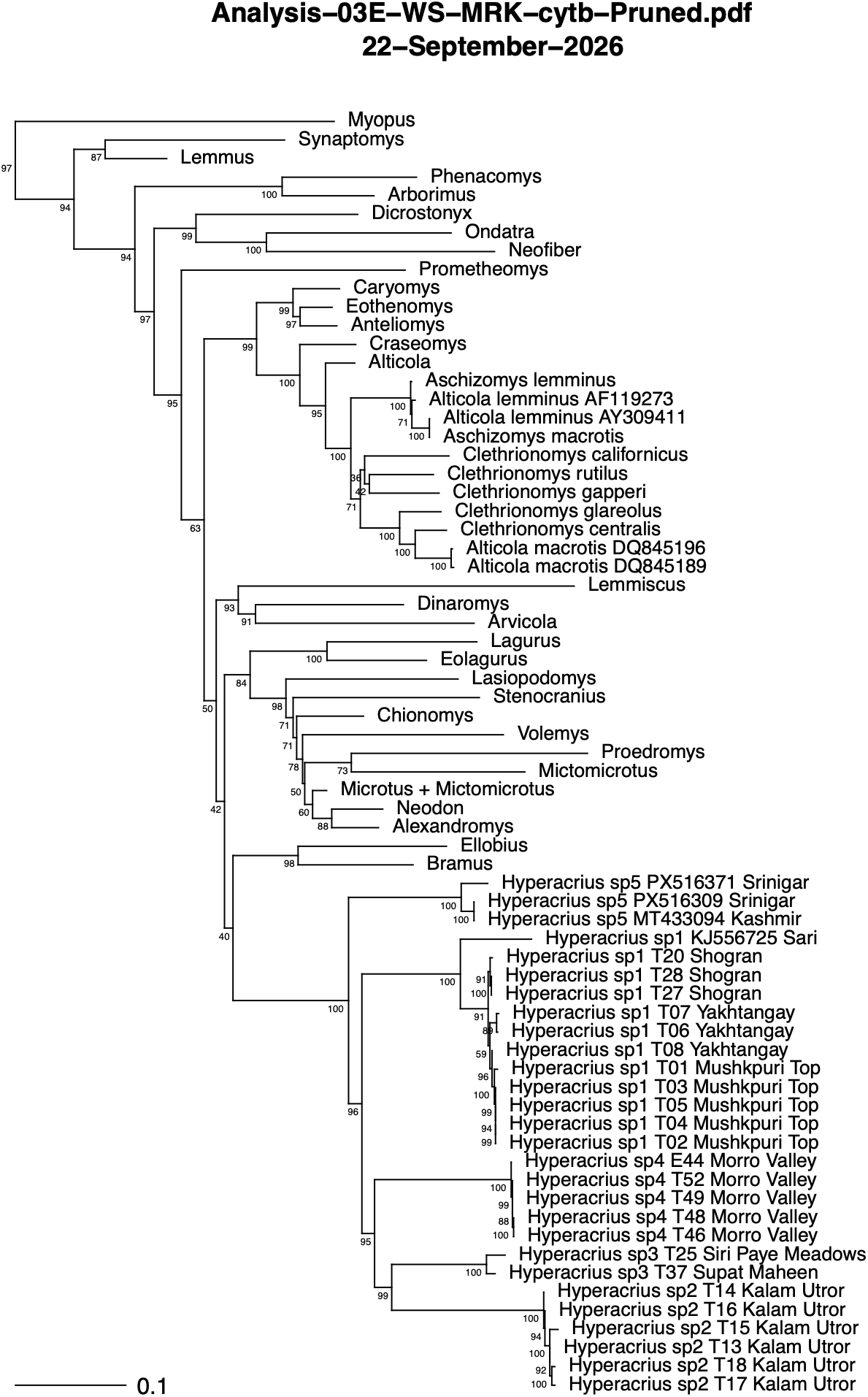
Maximum likelihood phylogeny using cyt b (Analysis 3). Genera with multiple species represented are pruned to the genus name. Ultrafast bootstrap values are shown at the nodes. The scale units are substitutions per site. The complete phylogeny with 194 tips is in Supplementary Data SD14.

Analysis 4. COI only (Fig. 9 and Supplementary Data SD15). This analysis has no data for *Hyperacrius*, and the topology is the most divergent. Microtini is a clade, but *Arvicola* is not the sister taxon. Between *Arvicola* and Microtini is a poorly supported group of genera that are usually recovered as early-diverging: *Lemmus*, *Synaptomys*, *Phenacomys*, *Ondatra*, as well as *Lagurus* and *Eolagurus*. The Clethrionomyini is the sister-group of all of these, although support for this clade is poor (ufbs = 45).

**Fig. 9.**
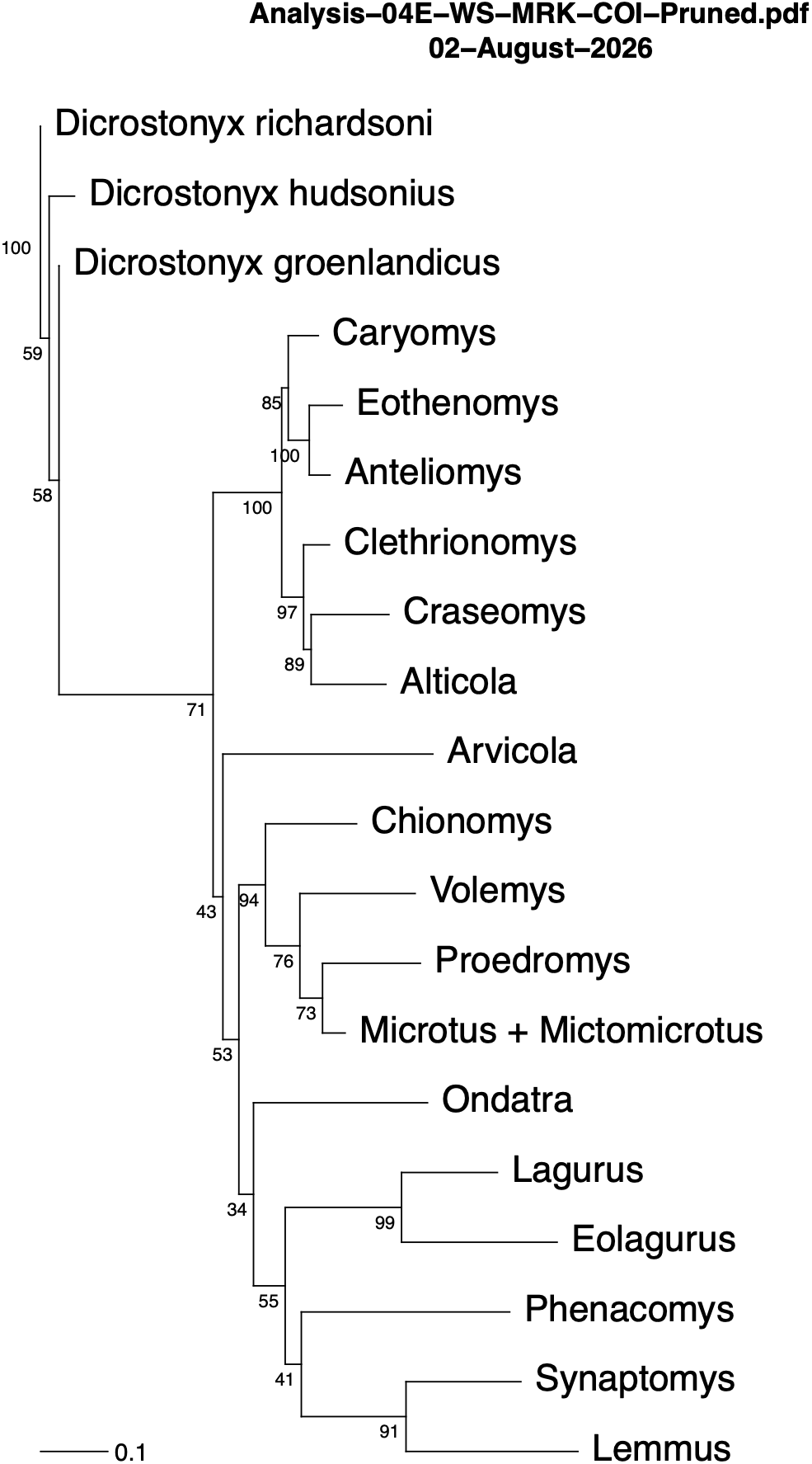
Maximum likelihood phylogeny using COI (Analysis 4). Genera with multiple species represented are pruned to the genus name. Ultrafast bootstrap values are shown at the nodes. The scale units are substitutions per site. The complete phylogeny with 194 tips is in Supplementary Data SD15.

Analysis 5. Cytb and three nuclear genes. (Fig. 10, Supplementary Data SD16). The topology is almost identical to that of Analysis 1, with similar bootstrap support values that support the placement of *Hyperacrius* as the sister group of the poorly supported clade described in Analysis 1.

**Fig. 10.**
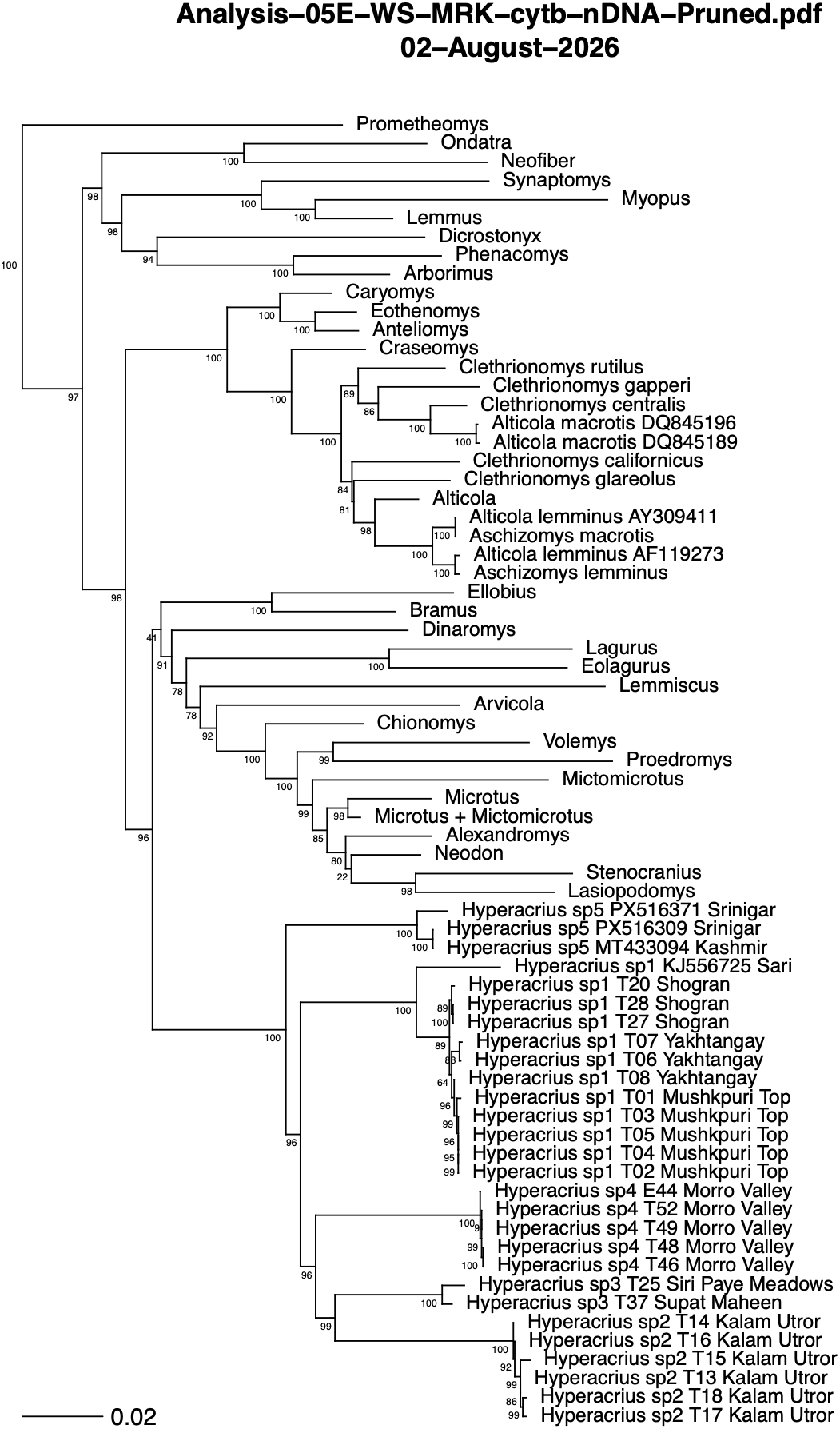
Maximum likelihood phylogeny using cyt b and nuclear genes (Analysis 5). Genera with multiple species represented are pruned to the genus name. Ultrafast bootstrap values are shown at the nodes. The scale units are substitutions per site. The complete phylogeny with 194 tips is in Supplementary Data SD16.

### Relationships of *Alticola* to other Clethrionomyini

Our sequences of *Alticola* were almost identical (<1% divergence in Cytb) to GenBank sequence KJ556728, *Alticola montosa*, which was not used by Withnell and Scarpetta (2024). We consider them to be conspecific with this sequence. We added KJ556728 to the Withnell-Scarpetta matrix, as well as GenBank sequences of *A. lemminus* (AY309411, AF119273), *A. macrotis* (DQ845196, DQ845189), and *A. stracheyi* (KY968269).

In all analyses (except 4) we found a clade of sequences of *Alticola montosa* and GenBank KJ556728 (*A. montosa*) to be the sister-group of *Alticola argentatus* (Supplementary Data SD12–16). In general, we found the same arrangement within Clethrionomyini as Withnell and Scarpetta (2024): (1) *Caryomys* + *Eothenomys* + *Anteliomys*, and (2) *Craseomys* + *Clethrionomys* + *Alticola*. *Alticola* was paraphyletic with respect to *Clethrionomys*, with *A*. *lemminus* and *A. macrotis* forming a clade more closely related to *Clethrionomys* than to other *Alticola*; alternatively, one could consider *Alticola* to be polyphyletic, based on the topology alone.

In Analysis 3 (Cytb only), *A. macrotis* (DQ845189 and DQ845196) is nested within *Clethrionomys*, and is the sister group to *C. centralis* (100% ufbs). The clade of three *Alticola lemminus* (AF119273, AY309411, and KJ556633) is strongly supported (ufbs = 100) as the sister-group of the clade containing *C. californicus*, *C. gappe*ri, *C. rutilis*, *C. glareolus*, and *C. centralis*.

In all analyses except for Analysis 4 we found discordance among species of *Clethrionomys*, *Alticola lemminus* and *A*. *macrotis*. *Clethrionomys* was paraphyletic with respect to *A. macrotis*, and in Analysis 5 *Clethrionomys* was also paraphyletic with respect to *Alticola lemminus*. We note that the *A. macrotis* sequence by Withnell and Scarpetta is likely to be *A. lemminus*. In all analyses, the two “*A. macrotis*” (DQ845196 and DQ845189) were nested within *Clethrionomys*. These two sequences, labelled as *A. macrotis vinogradovi* in GenBank, were sequenced by Lebedev et al. (2007). This incongruence (the close relationship of *A. macrotis* [DQ845196 and DQ845189] with *Clethrionomys centralis)* may represent ancient mitochondrial capture, but nuclear genes of *A. macrotis* need to be analyzed (Kohli et al. 2014).

## Discussion

### Species delimitation and systematics in *Hyperacrius*

The species-level systematics of *Hyperacrius* remains poorly studied, except for Phillips (1969). Kryštufek and Shenbrot (2022) summarized information about *Hyperacrius*. The species are found in the southern Himalayan mountains in Pakistan and India, inhabiting alpine meadows, forests, and grasslands from 1951–3658 m elevation. The Srinagar sequences (1,610 m per GenBank) would extend the lower limit, if the locality data are accurate).

Our study is based on a limited sample of *Hyperacrius* from seven populations, for which we sequenced Cytb, the gene most widely used in mammalian systematics. The five clades of *Hyperacrius* recovered are each well supported and are separated by large divergences. Mean uncorrected p-distances between clades are 11.9–13.8%, and the smallest pairwise distance between any two clades is 11.2% (Table 1). In contrast, mean distances within clades are 0.03–1.2%, and the maximum is 2.4%, meaning a gap of roughly an order of magnitude between intra- and interclade divergence.

Based on these distances we have treated each clade as a full species, cognizant of the fact that our primary data derive from one mitochondrial gene. However, these distances exceed proposed mammalian benchmarks for species status for Cytb. Bradley and Baker (2001) proposed that Kimura two-parameter distances below 2% indicate intraspecific variation and those above 11% indicate distinct species. Among the rodents they examined, mean distances were 0.85% within subspecies, 2.09% within species, and 9.55% between sister species. For *Microtus*, the arvicoline most comparable to *Hyperacrius*, sister species differ by 4.3–11.1% (mean 6.8%) (Baker and Bradley 2006:Table 3).

For comparison, we also calculated the K2P distances for *Hyperacrius* (Table 1). Mean K2P distances between clades are 13.3–15.7%, and the smallest pairwise distance between any two clades is 12.4% (Table 1). In contrast, mean distances within clades are 0.02–1.2%, and the maximum is 2.4%. Every interclade comparison in *Hyperacrius* exceeds the K2P threshold of Bradley and Baker (2001).

We do not regard the conclusions of Bradley and Baker as rules by which to delimit species, but rather as comparative evidence of separation, given that our distance data are comparable with interspecific distances between other rodent species. Reproductive isolation, reciprocal monophyly, or some degree of difference is evidence of separation of lineages or populations rather than a defining property of the species (de Queiroz 2020, 2025). Subspecies, whether defined traditionally as geographic subgroups that intergrade without evidence of reproductive isolation (Hillis 2021) or as incompletely separated lineages within a more inclusive lineage (de Queiroz 2020), require evidence of contemporary genetic exchange. Although our sample is small, we see no evidence of this, because no clades share haplotypes. Our K2P divergences of 13–16% imply isolation over a period far longer than that separating typical mammalian subspecies (Bradley and Baker 2001; Baker and Bradley 2006). Not surprisingly, most of the clades are known from valleys isolated by high ridges.

We also acknowledge that a critical test (*sensu* Wüster 2025) for distinct species should include large numbers of nuclear genes, ideally with multispecies coalescent analyses. Mitochondrial DNA alone cannot detect admixture, as one might expect in the case of subspecies, and it is subject to processes such as introgression and mitochondrial capture that may mislead inference. Several reptile species recognized or described primarily from mtDNA have been considered to be invalid (Wüster 2025), but these cases mostly involve widespread, geographically variable taxa with low genetic divergence and broad contact zones, in contrast to our *Hyperacrius* samples. Wüster (2025) also noted, however, that where candidate species are isolated by allopatry and differ by very high levels of mtDNA divergence, a confirmatory approach is likely to be adequate. Such is the case with *Hyperacrius*.

### Hyperacrius sp. 1: *Hyperacrius wynnei*

The type-locality of *Hyperacrius wynnei wynnei* is “Ad Mari (Murree) in montibus Himalayanis occidentalibus, ad latus occidentale fluminis Jhelum”, subsequently interpreted as “Murree, 2,135 m [7000 ft], North Punjab (border of Rawalpindi-Hazara districts) West Pakistan” (Phillips 1969:462) (Table 2). It is worth noting that “Ad Mari (Murree)” means “near Mari (Murree)”, not necessarily at the exact location of Murree. Phillips (1969) noted that the city of Murree is too low in elevation to be the type-locality, and therefore it is more likely that the specimens came from the nearby Murree Hills. However, the elevation of Murree is 2,291 m (7,516 ft), which is perhaps not too low for *Hyperacrius*.

Mushkpuri Top is only 17 km north of Murree (Fig. 1, Table 2) and given the low genetic divergence of Mushkpuri Top samples from other samples of *Hyperacrius* sp. 1 (Sari, Yakhtangay, and Shogran), we refer *H*. sp. 1 to *H. wynnei*.

Phillips (1969) named *Hyperacrius wynnei traubi* from “Yakh Tangai, 6600 ft, Swat, West Pakistan.” He stated that it was known only from the area of the type-locality but probably occurs in much of the southern part of Swat, below 8200 feet. He diagnosed *H. w. traubi* as having a longer zygomatic breadth and occipitonasal length than populations east of the Indus River, as well as different pelage coloration (Phillips 1969: Fig. 4 and Table 1).

Our samples from Yakhtangay, west of the Indus, differ from other samples of *H.* sp. 1 east of the river, including samples from near Murree, the type-locality of *H. wynnei wynnei*, by <1.4% p-distance and are genetically indistinguishable from them (Figs. 1, 2). We therefore consider them conspecific. If so, then *H. w. traubi* is a junior subjective synonym of *H. wynnei*, although we have not examined the pelage and skull characters that Phillips (1969) used to diagnose the subspecies. More broadly, the Indus River appears to have little effect on genetic differentiation within *H. wynnei*, a point also made for amphibians by Jablonski et al. (2026).

### Hyperacrius fertilis fertilis

We do not have any samples that we can refer to *Hyperacrius fertilis fertilis*. The location of the type-locality, “Pir Panjal,” is imprecise (Fig. 1), but information about W. L. Abbott’s expeditions in Taylor (2017) provides some information. Taylor (2017: Fig. 3) showed that from August to October of 1891 Abbott apparently travelled along the length of the Pir Panjal range itself, moving southeast to northwest. Specimens collected during this period are listed as “Pir Panjal range”, whereas other specimens collected by Abbott typically have more detailed information.

In his description of *H. f. brachelix*, Miller (1899) compared the type specimen (from Nagmarg) with the type series of *H. f. fertilis*, including the type series from the Pir Panjal range (8500 ft), Krishnagunga Valley (7000 ft), Kaj Nag Mountains (8000 ft), and Central Kashmir (12,000). However, elevation of the Central Kashmir specimen at 12,000 ft is anomalous. the USNM catalogue lists this specimen 20146/35509 as *Arvicola montosa* (= *Alticola montosa)*, not *Hyperacrius*. Evidently True misidentified this specimen in his original description of *H. fertilis* and Miller perpetuated the error. The correct identification of this specimen as *Alticola* explains its unusual elevation of 12,000 ft, because *Alticola* is a high-elevation vole.

Phillips (1969) referred a specimen from the type series (USNM 59393, collected by W. L. Abbott, 23 April 1891) from the Kaj Naj (= Kaj Nag or Kazinag) mountains (8000 ft) to *H. f. brachelix*. We view this identification as unlikely because the Kaj Nag mountains are considered part of the Pir Panjal range (although they lie north of the Jhelum River), which he claimed was within the distributional range of *H. fertilis fertilis.* Phillips (1969: Fig. 1) did not plot the location of Kaj Nag which suggests he did not know its location. We have labelled this small range on Fig. 1.

We follow Phillips (1969) and others in considering *Hyperacrius aitchisoni* (Fig. 1) to be a junior subjective synonym of *H. f. fertilis*.

### Hyperacrius sp. 5: Hyperacrius fertilis brachelix

We tentatively refer *Hyperacrius* sp. 5 to the nominal species *H. fertilis brachelix*, which we treat as a distinct species *H. brachelix.* The holotype was collected in November 1895 during W. L. Abbott’s third expedition, which was based out of Srinagar (Taylor 2017:Fig. 5). The species was described by Miller (1899). The type-locality is given as “Nagmarg, Kashmir (altitude, 9,000 feet)” (Poole and Schantz 1942). See Figure 1, locality P08.

We have no samples from the Kashmir Valley. However, GenBank accession MT433094 is from a partial mitochondrial genome from a specimen from “Kashmir”. According to Abramson et al. (2020), this voucher specimen (in the Zoological Institute, Russian Academy of Sciences) for MT433094 is labeled as *Hyperacrius fertilis brachelix* No. 29256, collector No. 76; collected 3.09.1903; Kashmir, 2,133.6 m. The specimen was provided in 1937 as a gift from the British Museum of Natural History.

The locality “Kashmir” is very common in museum records from the later 1800s and early 1900s, when the political lines for northwestern India and Pakistan were frequently redrawn. Until the mid-19th century, the term Kashmir included only the Kashmir Valley between the Great Himalayas and the Pir Panjal Range. However, the term has come to include a larger region that includes the princely state of Jammu and Kashmir (Fig. 1).

With a couple of assumptions, it is possible to link this specimen and therefore the DNA sequence to a nominal taxon. Colonel A. E. Ward collected specimens from “Kashmir” in the early 1900s that are currently in the American Museum of Natural History (AMNH) and the Natural History Museum, London (BMNH). VertNet records of AMNH specimens collected by A. E. Ward include Dachyani, Liddar Valley, Pahalgam, and Tullian. BMNH (Natural History Museum 2026a,b) records of *Hyperacrius* collected by Ward include Aru, Liddar Valley, Pahalgam, and Aston Marg. These localities are plotted on Fig. 1 as W01 to W06, and they are in the mountains forming the eastern rim of the Kashmir valley, east of Srinagar, the base for Ward’s expeditions. Based on the BMNH records, the number of *Hyperacrius* collected by Ward during 1902–1906 suggests that one of these specimens, collected in 1903 (“No. 29256, collector No. 76; collected 3.09.1903; Kashmir, 2,133.6 m”), was donated to the Russian Academy of Sciences and was used as the basis for the mitochondrial genome sequence MT433094.

The airline distance between Srinagar and Nagmarg (the type-locality of *H. brachelix*) is about 60 km. Based on the close sequence similarity between the Srinagar sequences (PX516309 and PX516371) and the mitogenome sequence (MT433094), which is likely from the mountains around Srinagar based on Ward’s collection sites, we tentatively link *Hyperacrius* sp. 5 to this nominal taxon and based on the level of sequence divergence from other *Hyperacrius* we recognize it as a full species, *Hyperacrius brachelix*.

Our inference calls into question the identification by Phillips (1969) of several specimens as *H*. *fertilis brachelix*. He examined specimens from Shogran (USNM 353587–88, elevation 10,300 ft) and Naran (Soch Valley) that he referred to *H. f. brachelix* and other specimens from Shogran that he referred to *H. wynnei wynnei* (USNM 353589–614, 7750 ft). Although Phillips (1969:469) referred specimens from the high elevations in the Kaghan Valley (e.g., Gitidas at 12,000 ft (= 3658 m)) to *H. f. brachelix*, the small elevation difference between *H*. sp. 3 (Supat Maheen, 3618 m) and Phillips *“brachelix”* suggests that the latter specimens might be referable *H.* sp. 3 rather than to *H. brachelix*. A detailed analysis of existing museum material, combined with DNA sampling of specimens will be necessary.

### Hyperacrius sp. 2: Hyperacrius fertilis zygomaticus

Given its proximity to the type-locality Utror (Fig. 1), *Hyperacrius* sp. 2 is referable to *Hyperacrius fertilis zygomaticus.* The mean pairwise p-distance to its closest relative, *Hyperacrius* sp. 3, is 13.0% (Table 1). Based on this level of divergence, it seems clear that *Hyperacrius fertilis zygomaticus* should be elevated to species status as *Hyperacrius zygomaticus*.

### *Hyperacrius* sp. 3: Supat Maheen and Siri Paye Meadows

It is possible that these population represent an undescribed species. *Hyperacrius* sp. 3 occurs at the highest elevations represented in our samples, at Supat Maheen (3618 m) and Siri Paye Meadows (3045 m). Phillips (1969: localities 16–17) reported examining *H. brachelix* from above tree line at Babusar (“9900 ft”) and Gitidas (“12,000 ft”), near the head of the Kaghan Valley. Given the proximity of these high-elevation localities to Supat Maheen and Siri Paye Meadows, one might conceivably refer *Hyperacrius* sp. 3 to *H. fertilis brachelix*. Indeed, specimens from Phillips’ (1969) localities 16, 17, and 18, all in the upper reaches of the Kaghan Valley (Fig. 1), may be conspecific with our *Hyperacrius* sp. 3.

However, most of the specimens that Phillips (1969) referred to *H. fertilis brachelix* are from localities near Nagmarg, the type-locality of *H. fertilis brachelix*, in the Kashmir Valley, and these specimens should be referable to *H. fertilis brachelix*. Nagmarg (Fig. 1, locality P08) is geographically distant from the upper Kaghan Valley. This geographic separation raises doubts about assigning the high-elevation Kaghan populations to *H. brachelix*. Consequently, *Hyperacrius* sp. 3 may not be referable to any nominal species.

### *Hyperacrius* sp. 4: Morro Valley

We consider it likely that this population represents an undescribed species. Some VertNet localities that are very near Morro Valley are relevant: Torbun, Lower Moru (2470 m); Lulu-Sar Lake (3048 m); Moru Top, Palas Valley, E of Pattan (2700 m), and examination of these specimens would be worthwhile.

### Morphometrics

#### Interspecific Analysis of Traits

In general, species at the high-elevation localities (Supat Maheen, Siri Paye Meadows) have shorter hind feet (HFL). Along with Morro Valley individuals, the Supat Maheen and Siri Paye Meadows voles also have relatively longer ears (EL and EL/HBL) (Fig. 3).

Phillips (1969:Table 1) found that skull measurements indicate that *H. wynnei* are slightly larger than *H. fertilis*. This agrees with our general result that specimens of *Hyperacrius* sp. 1 (*H. wynnei*: Shogran, Mushkpuri Top, and Yakhtangay) are larger than other *Hyperacrius*, with longer hindfeet (HFL) and shorter ear length (EL and EL/HBL). *Hyperacrius* sp. 2 differs in similar ways from *H.* sp. 3 and *H.* sp. 4 but it is not quite as different as is *H.* sp. 1. However, *Hyperacrius* sp. 2 has significantly shorter hind feet than *H.* sp. 1, but not when roughly corrected for body size (HFL/HBL).

### PCA Discussion

The overall pattern in the principal components analysis (Fig. 4) is best explained by between-group structure within the entire pooled sample. In a single homogeneous population, external measurements typically covary positively and PC1 reflects isometric size; negative cross-correlations of the kind implied by these loadings arise when groups that differ in proportion are analyzed together. The combined sample comprises several localities and species, and the opposition of body and foot dimensions against ear length on PC1 (Table 3) is consistent with voles from various sites, or of different species, differing in proportions rather than in size alone. Thus, PC1 is a group discriminant rather than a general size vector.

The appearance of a general size axis on PC2 rather than PC1 means that proportional differences between groups, even though these were not defined a priori, dominate the total variation; the general size axis is demoted to a secondary position. This is expected for a pooled sample of groups differing in external proportions.

### Linear Discriminant Analysis Discussion

The LDA results indicate that external body measurements distinguish *H.* sp. 1 clearly from the other three, largely on the basis of relatively short ears combined with proportionally longer hind feet and body length (Fig. 5, Table 4). *Hyperacrius* sp. 2 is moderately distinct but shows some morphometric overlap at both ends of its range, consistent with its intermediate classification accuracy.

In contrast, *H.* sp. 3 and *H.* sp. 4 are not well resolved by external measurements. They both have long ears, which separates them from *H.* sp. 1 and *H.* sp. 2, and the two are distinguished from each other by a secondary axis reflecting hindfoot and tail proportions. Not surprisingly, 4 of 10 *H.* sp. 4 individuals were misclassified as *H.* sp. 3 (Table 5), and the two species’ confidence ellipses overlap broadly. This suggests that hindfoot and tail length, the main contributors to LD2, do not reliably distinguish these two species.

Overall, the discriminant analysis is moderately consistent with the four-species classification scheme (70.3% overall accuracy versus a 25% chance baseline). The scheme lacks clear differentiation between *H*. sp. 3 and *H*. sp. 4 (Table 5).

### Phylogenetic relationships of *Hyperacrius* to other Arvicolinae

The phylogenetic position of *Hyperacrius* within Arvicolinae has long been uncertain. Morphology-based classifications assigned the genus either to Clethrionomyini (Hooper and Hart 1962; Gromov and Polyakov 1977) or to Lagurini (Kretzoi 1969). Miller (1896), followed by Phillips (1969), hypothesized that *Hyperacrius* was derived from *Alticola*, whereas Pozdnyakov (2008) left its tribal affiliation unresolved.

Early phylogenetic analyses disagreed about the position of *Hyperacrius*, and each had limited data or outgroup sampling. Robovský et al. (2008: Fig. 5), using a combined analysis of morphological and molecular data, recovered *Hyperacrius* near the base of Arvicolinae as the sister group of *Prometheomys*. They therefore regarded *Hyperacrius*, together with *Ellobius*, *Prometheomys*, and Lagurina, as representing basal arvicoline lineages. This placement is unusual, but it cannot be evaluated independently because their data and matrices are unavailable.

Kohli et al. (2014) published the first DNA sequences of *Hyperacrius*, in a molecular analysis of Clethrionomyini (formerly Myodini). Their Cytb analysis recovered *Hyperacrius* as sister to all other sampled Clethrionomyini with posterior probability >0.90 (their Fig. 2). However, their outgroups were limited to *Microtus* and *Dicrostonyx*, which makes the position of *Hyperacrius* difficult to assess. In their *G6pd* analysis, *Hyperacrius* fell in a polytomy that included *Microtus*, and the genus was not included in their multilocus species-tree analysis (their Fig. 3). They left its tribal placement unresolved. Similarly, a multilocus analysis by Tang et al. (2018: Fig. 3) recovered *Hyperacrius* as the sister taxon of Clethrionomyini (their Myodini) with strong support. Their only outgroups were *Microtus* and *Neodon*, however, so there is little confidence in the rooting.

Larger-scale analyses have produced conflicting results. The comprehensive synthesis of Upham et al. (2019) placed *Hyperacrius* nested within Clethrionomyini with strong support (their Supplementary Data S3; PP = 1.0). However, they included no sequence data for *Hyperacrius*, and the placement was based on imputation, even though sequences were apparently available from Kohli et al. (2014). In contrast, the analysis of mitochondrial protein-coding genes of Abramson et al. (2020: Fig. 3) placed *Hyperacrius* in Arvicolini, in a clade containing *Arvicola*, *Microtus*, and other microtine genera. Support for inclusion of *Hyperacrius* in this clade was strong (95–100), although support for its position as the sister-group of the other members was weak. Interpretation of this result is complicated by incomplete taxon sampling. Their analysis lacked several genera, including *Lemmiscus*, *Dinaromys*, *Bramus*, *Ellobius*, *Eolagurus*, and *Lagurus*, some of which occupy positions between Clethrionomyini and Arvicolini in other analyses with broader taxonomic sampling. Abramson et al. (2021: Fig. 2), with an expanded dataset, found *Hyperacrius* to be strongly supported as the sister group of the Microtini (including *Lemmiscus)*. However, the Arvicolini as delineated on their tree was not monophyletic, as Arvicola lay outside of the clade.

Kryštufek and Shenbrot (2022) named the subtribe Hyperacrina for *Hyperacrius* within an expanded tribe Arvicolini, which in their scheme includes the subtribes Pliomyina, Bramina, Lagurina, Arvicolina, and Microtina. That classification, which conflicts with that of Withnell and Scarpetta (2024), was restricted to Palearctic voles and lemmings. Moreover, proposing a new monotypic subtribe does not add to an understanding of the generic relationships.

As an aside, Hyperacrina is incorrectly formed: because the stem of *Hyperacrius* is *Hyperacri-*, the correct spelling is Hyperacriina. The Code (ICZN 1999: Articles 29.3, 32.5.3) requires correcting an incorrectly formed family-group name while keeping its authorship and date, so Hyperacriina Kryštufek and Shenbrot 2022 would be the correct citation and spelling.

Withnell and Scarpetta (2024) recovered *Hyperacrius* as the sister group of Clethrionomyini in their mtDNA maximum-likelihood and Bayesian phylogenies (their Appendices D and G), but with weak support. In their concatenated Bayesian analysis of nuclear and mitochondrial genes, however, *Hyperacrius* was the sister group of *Dinaromys*, a pliomyine.

Overall, the only set of relationships based on good taxon sampling and strong support for the position of *Hyperacrius* is that of Abramson et al. (2021), which places *Hyperacrius* closer to microtines than to Clethrionomyini.

In our analyses, Analysis 1 (Cytb, COI, and the nuclear genes) and Analysis 5 (Cytb and the nuclear genes) reveal support for the position of *Hyperacrius*. In both, *Hyperacrius* is the sister group of a poorly supported clade of the Microtini, Arvicolini, Ellobiusini, Lagurini, and *Dinaromys*. This clade + *Hyperacrius* is the sister-group of the Clethrionomyini, with very strong support (ufbs = 98).

This result is similar with that of Abramson et al. (2021). The two differ only in whether *Hyperacrius* is the sister group to Microtini alone or to a larger clade, and that larger clade is itself poorly supported. In contrast, analyses of Cytb alone (Analysis 3) and of Cytb + COI (Analysis 2) yielded poorly supported placements of *Hyperacrius*. Together, these results suggest that larger nuclear gene datasets may resolve intergeneric relationships in Arvicolinae. Cytb remains informative at shallow levels, as shown by its resolution of species within *Hyperacrius*. COI alone contributed little resolution to the topology overall, although adding it to Cytb and the nuclear genes did not reduce support (Analysis 1 vs. Analysis 5).

Previous and current analyses therefore indicate that *Hyperacrius* does not belong to Clethrionomyini, and hence that a close relationship with *Alticola*, as proposed by Phillips (1969), is unlikely. *Hyperacrius* belongs to a well-supported clade that also includes Clethrionomyini, Microtini, *Arvicola*, Lagurini, *Dinaromys*, and Ellobiusini. Within that clade it is more closely related to the Microtini group than to Clethrionomyini, but the lack of resolution among the lineages of the Microtini group continues to obscure its sister group.

### Monophyly and generic limits of *Alticola*

Musser and Carleton (2005) recognized three subgenera of *Alticola*. *Aschizomys* comprised *A. macrotis* and *A. lemminus*, *Platycranius* contained *A. strelzowi*, and the nominotypical subgenus *Alticola* included the remaining species. In our analyses, however, *A. strelzowi* is well nested within *Alticola*, so recognizing *Platycranius* would render the nominotypical subgenus paraphyletic. We therefore do not recognize *Platycranius*.

More broadly, our analyses (excluding 4 which has only one *Alticola*; Figs. 6–8 and 9) consistently recover *Alticola* as nonmonophyletic, in agreement with previous studies that found the genus to be paraphyletic or polyphyletic (Bodrov et al. 2020; Kryštufek and Shenbrot 2022; Dvoyashov et al. 2023). Sequences assigned to *Alticola* fall into three separate clades: (1) GenBank samples identified as *A. macrotis* (DQ845196 and DQ845189), nested within a clade of *Clethrionomys*; (2) *A. lemminus* together with other sequences identified as *A. macrotis*; and (3) the remaining species of *Alticola*.

This nonmonophyly may result, at least in part, from mitonuclear discordance. Using both mitochondrial and nuclear genes, Bodrov et al. (2020) inferred that *A. macrotis* had captured the mtDNA of *A. lemminus*. Their nuclear BRCA and GHR gene trees each recovered *A. lemminus* and *A. macrotis* as a monophyletic group (their Figs. S1, S3). However, their taxon sampling omitted several species of *Clethrionomys* that are important for evaluating the broader relationships of these taxa, including *C. californicus* and *C. gapperi* (their Fig. 3).

Resolving the nonmonophyly of *Alticola*, particularly the relationships and generic placement of *A. lemminus* and *A. macrotis*, is beyond the scope of this study. Two classifications are consistent with our results. First, a single genus could encompass *Clethrionomys*, *Alticola*, and *Aschizomys*. Because *Clethrionomys* Tilesius 1850 has priority over *Alticola* Blanford 1881 and *Aschizomys* Miller 1899, *Clethrionomys* would be the valid name for the combined genus (although it might be argued that *Myodes* Pallas 1811 has priority). Alternatively, *Alticola*, *Clethrionomys*, and *Aschizomys* could be recognized as distinct genera within Clethrionomyini and Hart, 1962, although the appropriate generic allocation of *A. macrotis* and *A. lemminus* remains uncertain. In either case, the type species of *Alticola* is *Arvicola stoliczkanus* Blanford 1875, so the name *Alticola* must remain with the clade containing that species. Consequently, a classification recognizing only *Alticola* and *Clethrionomys* would be difficult to reconcile with the relationships recovered here without substantially expanding the circumscription of one of those genera.

## Conclusions

Our results indicate that *Hyperacrius* is considerably more diverse than the two species recognized by Phillips (1969). Mitochondrial divergence of roughly 12–14% among five well-supported clades suggests that the Himalayan and Hindu Kush valleys of northern Pakistan and Kashmir harbor several deeply divergent, geographically restricted lineages, consistent with the high endemism expected in isolated mountain systems. The biogeographic patterns are not simple. The Indus River, regarded as a barrier separating *Hyperacrius* taxa (Phillips 1969), has apparently had little effect on genetic differentiation within *H. wynnei*. In contrast, distinct lineages replace one another over short distances within a single valley system. Elevation, local topography, and major ranges such as the Pir Panjal, which separates the Kashmir Valley from the Kaghan Valley, may therefore structure diversity in these voles more strongly than large rivers.

We recognize *H. wynnei* (with *H. w. traubi* as a junior subjective synonym), *H. zygomaticus*, new rank, and, tentatively, *H. brachelix*. Two further lineages, from the upper Kaghan Valley and the Morro Valley, may represent undescribed species. External measurements corroborate some of these distinctions but not others, so body proportions alone are insufficient for identification, particularly for the high-elevation lineages.

At higher levels, our analyses reject the traditional placement of *Hyperacrius* in Clethrionomyini, but its position among the remaining arvicoline lineages remains unresolved. Within Clethrionomyini, the nonmonophyly of *Alticola* and the nesting of *A. macrotis* sequences within *Clethrionomys* indicate that generic limits in this group also require revision.

Several lines of research would resolve the questions raised here. Our new data were limited to a single gene, albeit the one most commonly used in rodent systematics. Given that our first goal was to sort out relationships among individual *Hyperacrius*, and second, to infer the relationships of *Hyperacrius* to other arvicolines, this was appropriate. However, nuclear and genome-scale data from multiple individuals are needed to critically test the mitochondrial species boundaries as well as to place *Hyperacrius* properly within Arvicolinae.

Sampling is needed from the type localities and historical collecting areas, especially the Pir Panjal range, for the apparently unsampled *H. f. fertilis*; Nagmarg and the eastern rim of the Kashmir Valley, for *H. brachelix*; and the Indian portion of the range.

DNA from museum specimens, particularly those collected by W. L. Abbott and A. E. Ward and the material examined by Phillips (1969), could link nominal taxa to genetic clades. Also, description of the upper Kaghan and Morro Valley lineages as species will require cranial and dental morphometrics in addition to external measurements. Our morphometric analysis was limited to small samples of the specimens of four species that we collected, and we were not able to distinguish all species. We also did not examine the extensive *Hyperacrius* collections at the USNM or the University of Florida.

Finally, we offer that single-gene assessments remain valuable even in the face of the ability to produce gigabases of genomic sequence data. For groups such as *Hyperacrius* that are in dark areas of the Tree of Life, we agree with recommendations to describe species on limited evidence rather than waiting for comprehensive sampling (Hillis 2019). We therefore regard our samples as representing five well-supported species, and welcome further testing with genomic data and denser sampling of voles.

## Supporting information

Supp Mat 1

Supp Mat 2

Supp Mat 3

Supp Mat 4

Supp Mat 5

Supp Mat 6

Supp Mat 7

Supp Mat 8

Supp Mat 10

Supp Mat 11

Supp Mat 12

Supp Mat 13

Supp Mat 14

Supp Mat 15

Supp Mat 16

## Acknowledgments

We thank Kelly Zamudio and Anat Belasen for access to lab space and assistance with techniques. The Khyber Pakhtunkhwa wildlife department and local people assisted in fieldwork.

## Author Contributions

M. Raza Khan (Conceptualization, Field work, Lab Work, Data Analysis, Writing—original draft), Jeff Coleman (Lab Work, Writing—review and editing), M. Sajid Nadeem (Supervision, Conceptualization, Writing—review and editing), and David Cannatella (Conceptualization, Supervision, Funding, Software, Data Analysis, Project Administration, Writing—review and editing)

## Data Availability

All data, code, and analysis results are available in Supplementary Data files.

## Supplementary Data

Supplementary Data SD1.—Georeferenced localities from Phillips (1969) and collecting localities of A. E. Ward.

Supplementary Data SD2.—Classification of genera by tribe, following Withnell and Scarpetta (2024).

Supplementary Data SD3.—Nexus file of sequences; 194 tips, five genes.

Supplementary Data SD4.—Corrections to Withnell and Scarpetta (2024) matrix.

Supplementary Data SD5.—Complete edited table of WS-MRK sequences.

Supplementary Data SD6.—External measurements of *Hyperacrius* specimens used in morphometric analyses.

Supplementary Data SD7.—R scripts for morphometric analysis.

Supplementary Data SD8.—Maximum likelihood parameters of phylogenetic analyses.

Supplementary Data SD9.—Results for Kruskal–Wallis and Dunn tests performed on species groups.

Supplementary Data SD10.—Tables of diagnostics from the principal components analysis

Supplementary Data SD11.—Tables of diagnostics from the linear discriminant analysis with a priori species groups.

Supplementary Data SD12.—Phylogeny of five genes; log file of Analysis-01E (.zip).

Supplementary Data SD13.—Phylogeny of with mtDNA only genes; log file of Analysis-02E (.zip).

Supplementary Data SD14.—Phylogeny of Cytb only; log file of Analysis-03E (.zip).

Supplementary Data SD15.—Phylogeny of COI only; log file of Analysis-04E (.zip).

Supplementary Data SD16.—Phylogeny of Cytb and three nuclear genes; log file of Analysis-05E (.zip).

## Appendix Tissue samples sequenced

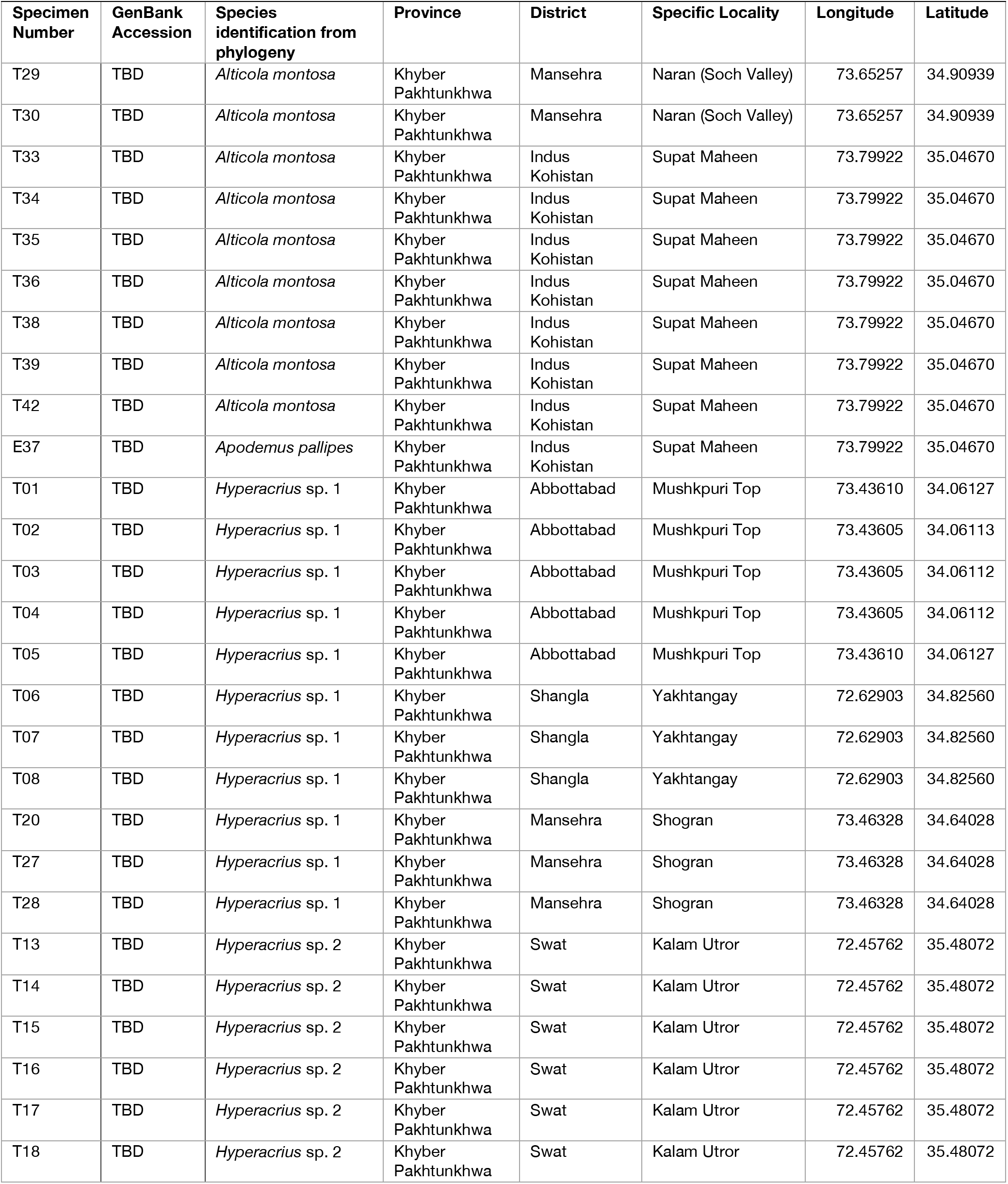
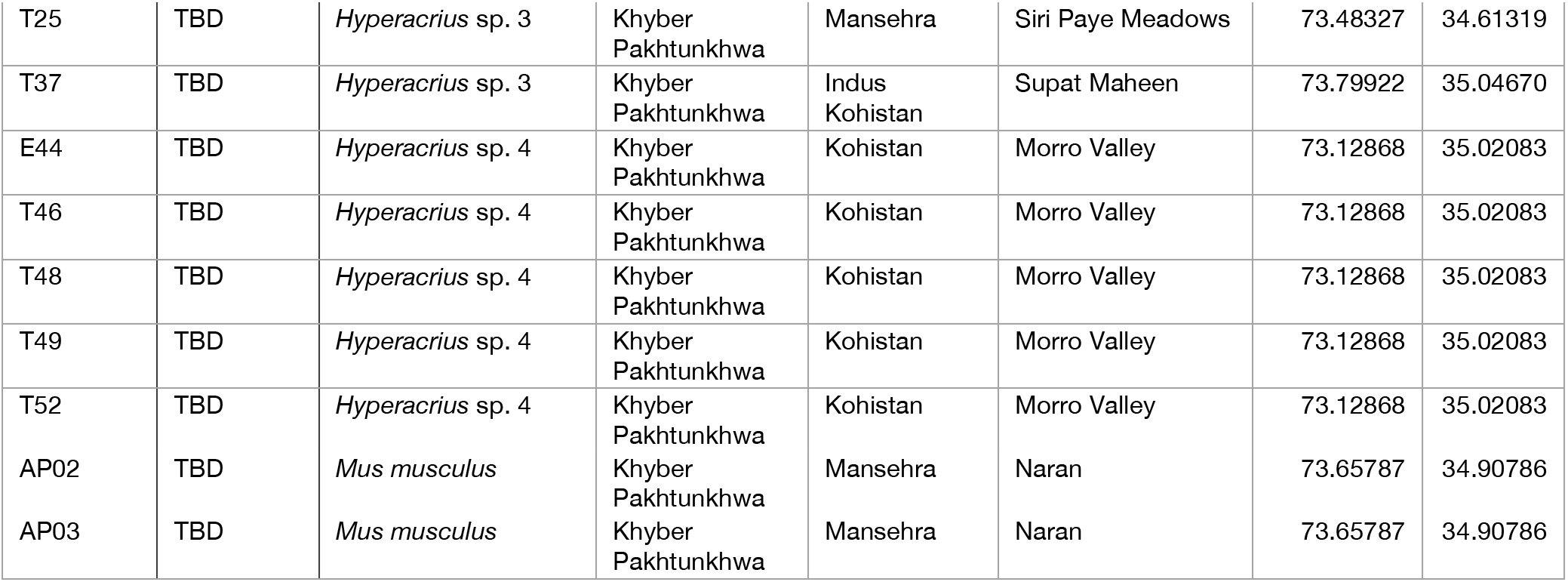

| Specimen Number | GenBank Accession | Species identification from phylogeny | Province | District | Specific Locality | Longitude | Latitude |
| --- | --- | --- | --- | --- | --- | --- | --- |
| T29 | TBD | *Alticola montosa* | Khyber Pakhtunkhwa | Mansehra | Naran (Soch Valley) | 73.65257 | 34.90939 |
| T30 | TBD | *Alticola montosa* | Khyber Pakhtunkhwa | Mansehra | Naran (Soch Valley) | 73.65257 | 34.90939 |
| T33 | TBD | *Alticola montosa* | Khyber Pakhtunkhwa | Indus Kohistan | Supat Maheen | 73.79922 | 35.04670 |
| T34 | TBD | *Alticola montosa* | Khyber Pakhtunkhwa | Indus Kohistan | Supat Maheen | 73.79922 | 35.04670 |
| T35 | TBD | *Alticola montosa* | Khyber Pakhtunkhwa | Indus Kohistan | Supat Maheen | 73.79922 | 35.04670 |
| T36 | TBD | *Alticola montosa* | Khyber Pakhtunkhwa | Indus Kohistan | Supat Maheen | 73.79922 | 35.04670 |
| T38 | TBD | *Alticola montosa* | Khyber Pakhtunkhwa | Indus Kohistan | Supat Maheen | 73.79922 | 35.04670 |
| T39 | TBD | *Alticola montosa* | Khyber Pakhtunkhwa | Indus Kohistan | Supat Maheen | 73.79922 | 35.04670 |
| T42 | TBD | *Alticola montosa* | Khyber Pakhtunkhwa | Indus Kohistan | Supat Maheen | 73.79922 | 35.04670 |
| E37 | TBD | *Apodemus pallipes* | Khyber Pakhtunkhwa | Indus Kohistan | Supat Maheen | 73.79922 | 35.04670 |
| T01 | TBD | *Hyperacrius* sp. 1 | Khyber Pakhtunkhwa | Abbottabad | Mushkpuri Top | 73.43610 | 34.06127 |
| T02 | TBD | *Hyperacrius* sp. 1 | Khyber Pakhtunkhwa | Abbottabad | Mushkpuri Top | 73.43605 | 34.06113 |
| T03 | TBD | *Hyperacrius* sp. 1 | Khyber Pakhtunkhwa | Abbottabad | Mushkpuri Top | 73.43605 | 34.06112 |
| T04 | TBD | *Hyperacrius* sp. 1 | Khyber Pakhtunkhwa | Abbottabad | Mushkpuri Top | 73.43605 | 34.06112 |
| T05 | TBD | *Hyperacrius* sp. 1 | Khyber Pakhtunkhwa | Abbottabad | Mushkpuri Top | 73.43610 | 34.06127 |
| T06 | TBD | *Hyperacrius* sp. 1 | Khyber Pakhtunkhwa | Shangla | Yakhtangay | 72.62903 | 34.82560 |
| T07 | TBD | *Hyperacrius* sp. 1 | Khyber Pakhtunkhwa | Shangla | Yakhtangay | 72.62903 | 34.82560 |
| T08 | TBD | *Hyperacrius* sp. 1 | Khyber Pakhtunkhwa | Shangla | Yakhtangay | 72.62903 | 34.82560 |
| T20 | TBD | *Hyperacrius* sp. 1 | Khyber Pakhtunkhwa | Mansehra | Shogran | 73.46328 | 34.64028 |
| T27 | TBD | *Hyperacrius* sp. 1 | Khyber Pakhtunkhwa | Mansehra | Shogran | 73.46328 | 34.64028 |
| T28 | TBD | *Hyperacrius* sp. 1 | Khyber Pakhtunkhwa | Mansehra | Shogran | 73.46328 | 34.64028 |
| T13 | TBD | *Hyperacrius* sp. 2 | Khyber Pakhtunkhwa | Swat | Kalam Utror | 72.45762 | 35.48072 |
| T14 | TBD | *Hyperacrius* sp. 2 | Khyber Pakhtunkhwa | Swat | Kalam Utror | 72.45762 | 35.48072 |
| T15 | TBD | *Hyperacrius* sp. 2 | Khyber Pakhtunkhwa | Swat | Kalam Utror | 72.45762 | 35.48072 |
| T16 | TBD | *Hyperacrius* sp. 2 | Khyber Pakhtunkhwa | Swat | Kalam Utror | 72.45762 | 35.48072 |
| T17 | TBD | *Hyperacrius* sp. 2 | Khyber Pakhtunkhwa | Swat | Kalam Utror | 72.45762 | 35.48072 |
| T18 | TBD | *Hyperacrius* sp. 2 | Khyber Pakhtunkhwa | Swat | Kalam Utror | 72.45762 | 35.48072 |
| T25 | TBD | *Hyperacrius* sp. 3 | Khyber<br>Pakhtunkhwa | Mansehra | Siri Paye Meadows | 73.48327 | 34.61319 |
| T37 | TBD | *Hyperacrius* sp. 3 | Khyber<br>Pakhtunkhwa | Indus<br>Kohistan | Supat Maheen | 73.79922 | 35.04670 |
| E44 | TBD | *Hyperacrius* sp. 4 | Khyber<br>Pakhtunkhwa | Kohistan | Morro Valley | 73.12868 | 35.02083 |
| T46 | TBD | *Hyperacrius* sp. 4 | Khyber<br>Pakhtunkhwa | Kohistan | Morro Valley | 73.12868 | 35.02083 |
| T48 | TBD | *Hyperacrius* sp. 4 | Khyber<br>Pakhtunkhwa | Kohistan | Morro Valley | 73.12868 | 35.02083 |
| T49 | TBD | *Hyperacrius* sp. 4 | Khyber<br>Pakhtunkhwa | Kohistan | Morro Valley | 73.12868 | 35.02083 |
| T52 | TBD | *Hyperacrius* sp. 4 | Khyber<br>Pakhtunkhwa | Kohistan | Morro Valley | 73.12868 | 35.02083 |
| AP02 | TBD | *Mus musculus* | Khyber<br>Pakhtunkhwa | Mansehra | Naran | 73.65787 | 34.90786 |
| AP03 | TBD | *Mus musculus* | Khyber<br>Pakhtunkhwa | Mansehra | Naran | 73.65787 | 34.90786 |

