## Supplementary material for "Insights into the molecular systematics of montane voles of the Western Himalayan Mountains": Supp Mat 4

Supplementary Data SD2.—Corrections to Withnell and Scarpetta matrix

[This section OK]

All sequences used by Withnell and Scarpetta, as well as additional sequences were analyzed with BLAST to confirm species identity. Some corrections were made to the Withnell and Scarpetta (2024) matrix, and hereafter the matrix is referred to as WS.

Anomalous Microtus ochrogaster sequences.

Withnell and Scarpetta (2024) found discordant placements of *Microtus* *ochrogaster* in their trees. Their ML and bayesian analyses of mtDNA genes placed it outside of all other *Microtus* as the sister-group of all other Microtini (their Figs. 1A–B), whereas their analysis of nuclear genes only (their Figs. 2A–B) and combined genes (their Figs. 3A–B) placed it within *Microtus*, but with low support (see their Appendices D–I in Supplementary Materials).

A BLAST analysis revealed that the closest match to the *M. ochrogaster* COI sequence (JQ601444) is *Alexandromys* *kikuchii*, with 85.8% identity. Other close matches include *Neodon* *sikimensis* (85.4%) and *Chionomys* *roberti* (85.1%). This sequence is highly divergent (~85%) from the *M. ochrogaster* sequences JF456728–30 and KT166982, whereas the latter differ from each other at most by ~3%. This indicates that COI sequence JQ601444 is not from *Microtus* or likely any other microtine.

In contrast, BLAST analysis of the Cytb sequence matches other *M. ochrogaster* at >99% identity, suggesting correct identification. Because of the uncertainty in the COI sequence, we replaced both the Cytb and COI sequences used by WS with sequences from a mitochondrial genome (KT166982) of *M. ochrogaster.*

The provenance of the voucher specimens for JQ601444 (ROM 109826) and JF456728–30 (*M.* *ochrogaster*) is "Canada: Okotoks, Mercier's Property." These were sequenced as part of a barcode project by the Biodiversity Institute of Ontario. This large sequence divergence between JQ601444 and *Microtus* *ochrogaster* sequences and the fact that the sequence is not close to any other Genbank sequence suggests that it represents a sequencing error.

Other anomalous sequences.

Two other species were removed from the Withnell-Scarpetta matrix: *Synaptomys* *borealis* and *Volemys* *millicens*. For *Synaptomys* *borealis*, a BLAST search matched JF457125 (COI) blasted to *Ellobius* *tancrei* at 86.9%. However, several other GenBank sequences of *S. borealis* show identities with each other at 100% indicating that JF457125 is anomalous. Interestingly both JF457125 and JQ60144 were sequenced as part of a Canadian barcoding project headed by P. D. N. Hebert.

For *Volemys*, three sequences from different genes (JF906123, JF906135, and GQ374481) blasted to *Neodon* *chayunensis* with greater than >99.7 identity, so this species was removed.

We found an obvious misalignment of the Cytb and COI sequences in two *Craseomys* species in the WS "mitochondrial only.nex" matrix. However, this error was not found in the matrix of concatenated mitochondrial and nuclear genes ("combined.nex"), so that only the latter was used for analyses.
