## Supplementary material for "Insights into the molecular systematics of montane voles of the Western Himalayan Mountains": Supp 9

```
2026-08-01

Supplementary-Data-SD9-Kruskal-dunn-test-Species

--- Kruskal-Wallis test: TL ---
  H(3) = 10.038, p = 0.0182

--- Dunn test: TL ---
  Kruskal-Wallis rank sum test

data: x and group
Kruskal-Wallis chi-squared = 10.0383, df = 3, p-value = 0.02


                           Comparison of x by group
                             (Benjamini-Hochberg)
Col Mean-|
Row Mean |        sp1        sp2        sp3
---------+---------------------------------
     sp2 |   1.377739
         |     0.2524
         |
     sp3 |   2.894506   1.409562
         |    0.0228*     0.3173
         |
     sp4 |   2.427656   0.886573  -0.599235
         |    0.0456*     0.4504     0.5490

alpha = 0.05
Reject Ho if p <= alpha
--------------------------------------------------


--- Kruskal-Wallis test: HBL ---
  H(3) = 11.277, p = 0.0103

--- Dunn test: HBL ---
  Kruskal-Wallis rank sum test

data: x and group
Kruskal-Wallis chi-squared = 11.2769, df = 3, p-value = 0.01


                           Comparison of x by group
                             (Benjamini-Hochberg)
Col Mean-|
Row Mean |        sp1        sp2        sp3
---------+---------------------------------
     sp2 |   1.162649
         |     0.2940
         |
     sp3 |   2.961123   1.671358
         |    0.0184*     0.1893
         |
     sp4 |   2.639014   1.291961  -0.469804
         |    0.0249*     0.2946     0.6385

alpha = 0.05
Reject Ho if p <= alpha
--------------------------------------------------


--- Kruskal-Wallis test: TAIL ---
  H(3) = 5.479, p = 0.14

--- Dunn test: TAIL ---
  Kruskal-Wallis rank sum test

data: x and group
Kruskal-Wallis chi-squared = 5.4791, df = 3, p-value = 0.14


                           Comparison of x by group
                             (Benjamini-Hochberg)
Col Mean-|
Row Mean |        sp1        sp2        sp3
---------+---------------------------------
     sp2 |   1.558534
         |     0.2382
         |
     sp3 |  -0.795822  -2.187951
         |     0.6392     0.1720
         |
     sp4 |  -0.351106  -1.850139   0.456163
         |     0.7255     0.1929     0.7779

alpha = 0.05
Reject Ho if p <= alpha
--------------------------------------------------


--- Kruskal-Wallis test: HFL ---
  H(3) = 21.531, p = <0.001

--- Dunn test: HFL ---
  Kruskal-Wallis rank sum test

data: x and group
Kruskal-Wallis chi-squared = 21.5311, df = 3, p-value = 0


                           Comparison of x by group
                             (Benjamini-Hochberg)
Col Mean-|
Row Mean |        sp1        sp2        sp3
---------+---------------------------------
     sp2 |   3.044778
         |    0.0070*
         |
     sp3 |   4.456584   1.312019
         |    0.0000*     0.2274
         |
     sp4 |   2.275501  -0.886595  -2.269585
         |    0.0458*     0.3753    0.0348*

alpha = 0.05
Reject Ho if p <= alpha
--------------------------------------------------


--- Kruskal-Wallis test: EL ---
  H(3) = 23.902, p = <0.001

--- Dunn test: EL ---
  Kruskal-Wallis rank sum test

data: x and group
Kruskal-Wallis chi-squared = 23.9015, df = 3, p-value = 0


                           Comparison of x by group
                             (Benjamini-Hochberg)
Col Mean-|
Row Mean |        sp1        sp2        sp3
---------+---------------------------------
     sp2 |  -1.152852
         |     0.2988
         |
     sp3 |  -4.108472  -2.746717
         |    0.0002*    0.0120*
         |
     sp4 |  -3.837281  -2.405321   0.489972
         |    0.0004*    0.0242*     0.6242

alpha = 0.05
Reject Ho if p <= alpha
--------------------------------------------------


--- Kruskal-Wallis test: HFL/HBL ---
  H(3) = 5.509, p = 0.138

--- Dunn test: HFL/HBL ---
  Kruskal-Wallis rank sum test

data: x and group
Kruskal-Wallis chi-squared = 5.5092, df = 3, p-value = 0.14


                           Comparison of x by group
                             (Benjamini-Hochberg)
Col Mean-|
Row Mean |        sp1        sp2        sp3
---------+---------------------------------
     sp2 |   1.571519
         |     0.2321
         |
     sp3 |   1.085797  -0.451391
         |     0.4164     0.6517
         |
     sp4 |  -0.585660  -2.078914  -1.603106
         |     0.6697     0.2258     0.3267

alpha = 0.05
Reject Ho if p <= alpha
--------------------------------------------------


--- Kruskal-Wallis test: EL/HBL ---
  H(3) = 27.954, p = <0.001

--- Dunn test: EL/HBL ---
  Kruskal-Wallis rank sum test

data: x and group
Kruskal-Wallis chi-squared = 27.9535, df = 3, p-value = 0


                           Comparison of x by group
                             (Benjamini-Hochberg)
Col Mean-|
Row Mean |        sp1        sp2        sp3
---------+---------------------------------
     sp2 |  -1.280442
         |     0.2405
         |
     sp3 |  -4.549921  -3.038393
         |    0.0000*    0.0048*
         |
     sp4 |  -4.040157  -2.467211   0.735535
         |    0.0002*    0.0204*     0.4620

alpha = 0.05
Reject Ho if p <= alpha
--------------------------------------------------


--- Kruskal-Wallis test: TAIL/HBL ---
  H(3) = 10.631, p = 0.0139

--- Dunn test: TAIL/HBL ---
  Kruskal-Wallis rank sum test

data: x and group
Kruskal-Wallis chi-squared = 10.6309, df = 3, p-value = 0.01


                           Comparison of x by group
                             (Benjamini-Hochberg)
Col Mean-|
Row Mean |        sp1        sp2        sp3
---------+---------------------------------
     sp2 |   0.567967
         |     0.6841
         |
     sp3 |  -2.306945  -2.671715
         |    0.0632*    0.0453*
         |
     sp4 |  -1.868348  -2.277371   0.538864
         |     0.0926    0.0455*     0.5900

alpha = 0.05
Reject Ho if p <= alpha
--------------------------------------------------
```
