## Supplementary material for "Insights into the molecular systematics of montane voles of the Western Himalayan Mountains": Supp Mat 11

```
----LDA results/statistics (Species)------
2026-08-05

Supplementary-Data-SD11-Hyperacrius-lda-diagnostics-Species


--- Raw discriminant coefficients ---
         LD1     LD2     LD3
HBL  -0.0718 -0.0776 -0.0555
TAIL -0.1184  0.1242 -0.3489
HFL  -0.2998  0.9527  0.3848
EL    1.0719  0.4211  0.2109

--- Standardized discriminant coefficients ---
         LD1     LD2     LD3
HBL  -0.6653 -0.7194 -0.5141
TAIL -0.3247  0.3409 -0.9573
HFL  -0.3273  1.0401  0.4201
EL    1.0688  0.4199  0.2102

--- Structure coefficients (variable-LD correlations) ---
         LD1     LD2     LD3
HBL  -0.5858 -0.0466 -0.0866
TAIL  0.1781  0.4693 -0.8513
HFL  -0.7315  0.5827  0.2151
EL    0.8754  0.2363 -0.1445

--- Top contributor per axis ---
By |standardized coef|: LD1 = EL | LD2 = HFL
By |structure coef|:    LD1 = EL | LD2 = HFL

--- Proportion Trace ---
      LD1       LD2       LD3
84.134677 14.278060  1.587263

--- Leave-one-out CV confusion matrix ---
        Predicted
Observed sp1 sp2 sp3 sp4
     sp1  10   1   0   0
     sp2   2   5   0   1
     sp3   0   1   6   1
     sp4   1   0   4   5
LOO cross-validated accuracy: 70.3%

--- Per-Species correct-classification rate (LOO) ---
  sp1   sp2   sp3   sp4
0.909 0.625 0.750 0.500
```
