## Supplementary figures and images for "Insights into the molecular systematics of montane voles of the Western Himalayan Mountains"

### SD12-Analysis-01E-WS-MRK-All-Genes.pdf

#### Analysis−01E−WS−MRK−All−Genes.pdf
#### 01−August−2026

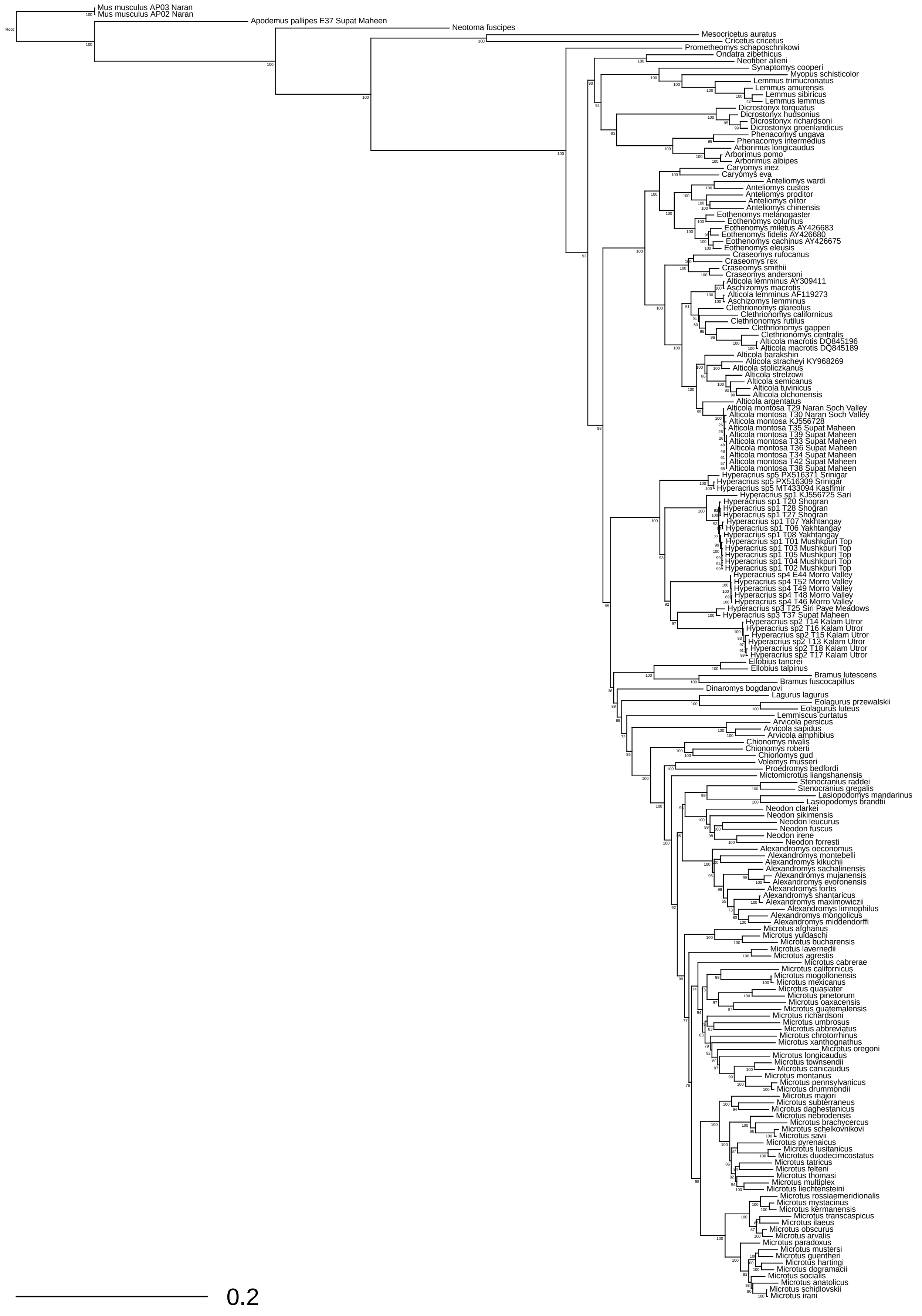

### SD13-Analysis-02E-WS-MRK-mtDNA.pdf

#### Analysis−02E−WS−MRK−mtDNA.pdf
#### 02−August−2026

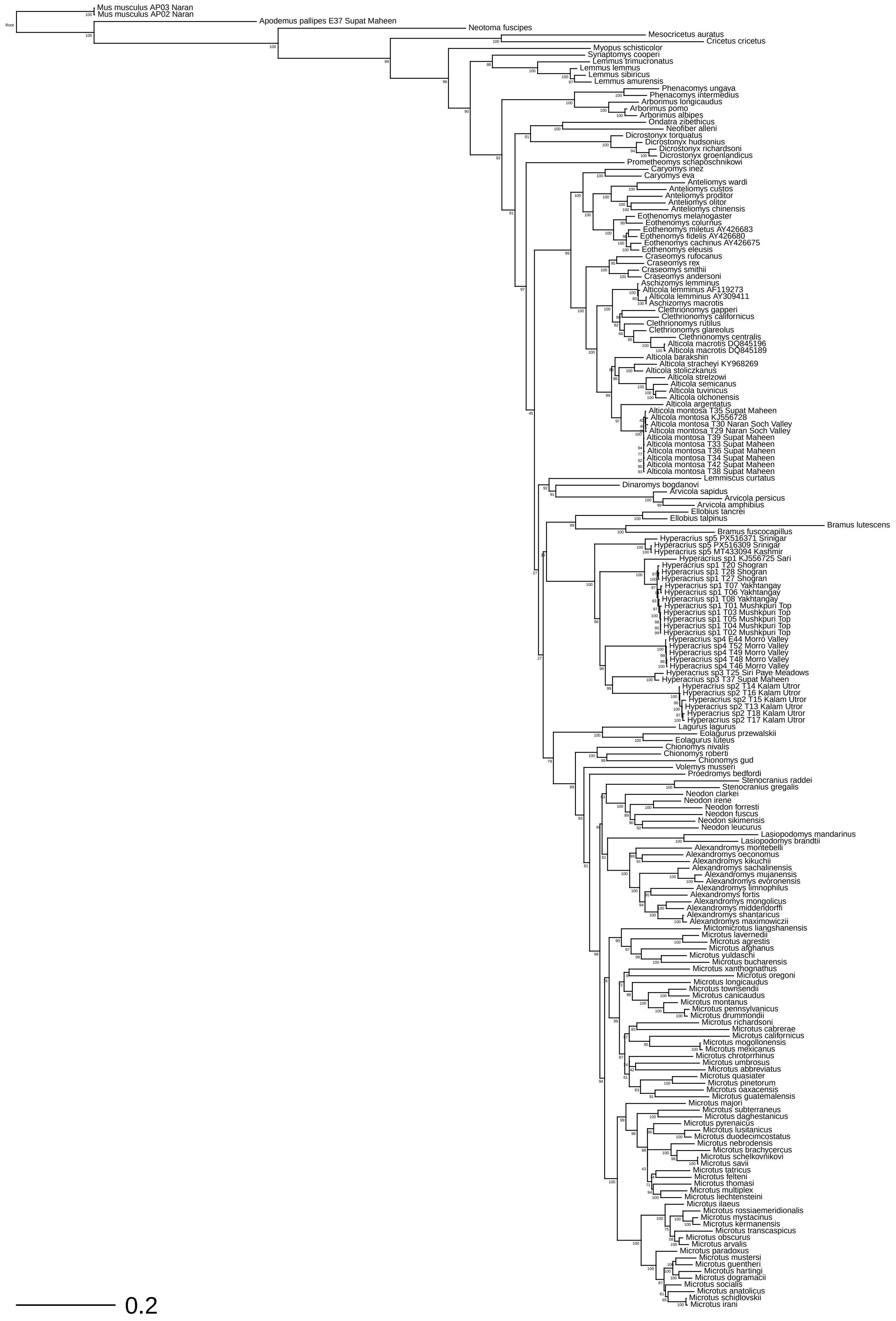

### SD14-Analysis-03E-WS-MRK-cytb.pdf

#### Analysis−03E−WS−MRK−cytb.pdf
#### 22−September−2026

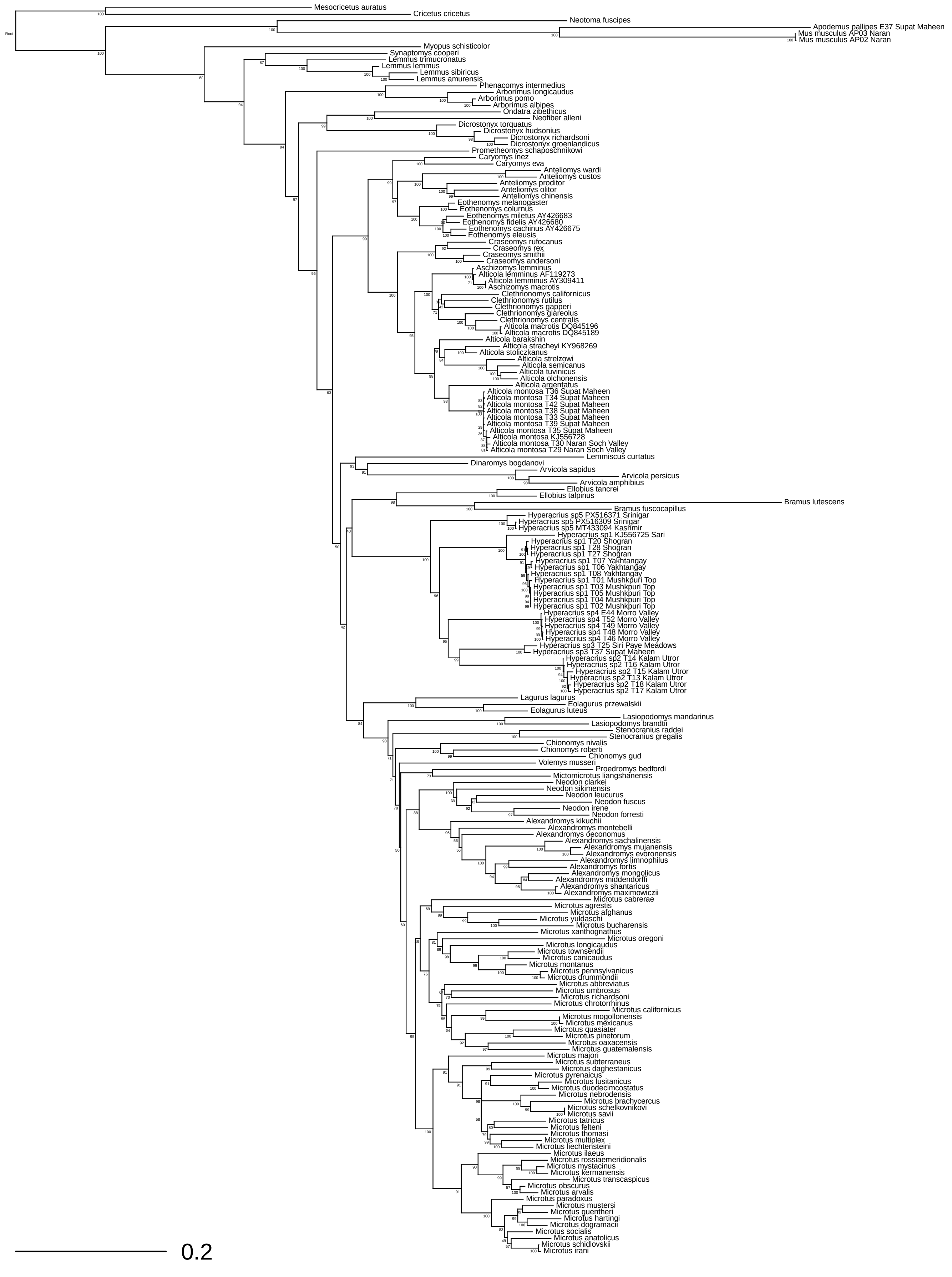

### SD15-Analysis-04E-WS-MRK-COI.pdf

#### Analysis−04E−WS−MRK−COI.pdf
#### 02−August−2026

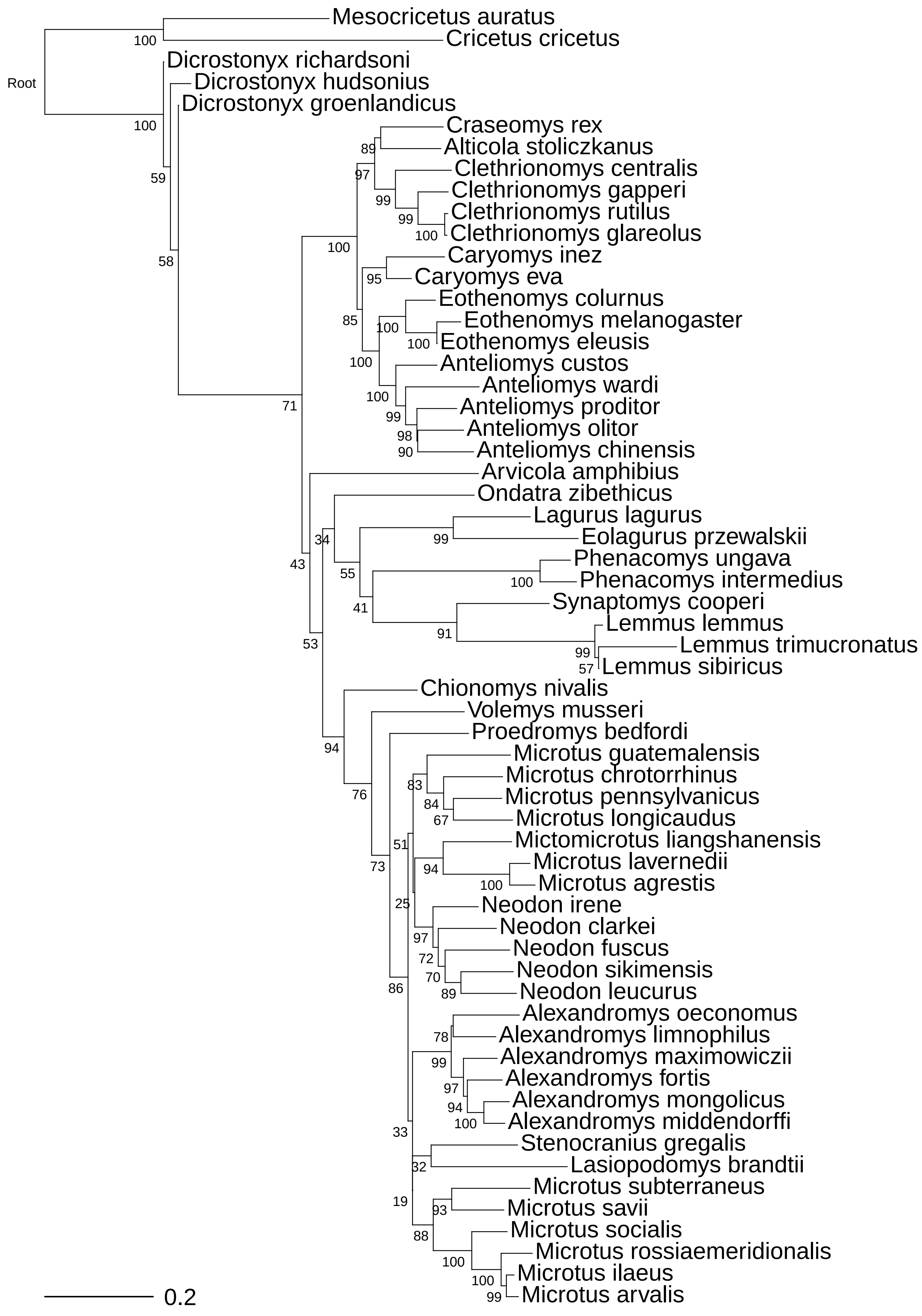

### SD16-Analysis-05E-WS-MRK-cytb-nDNA.pdf

#### Analysis−05E−WS−MRK−cytb−nDNA.pdf
#### 02−August−2026

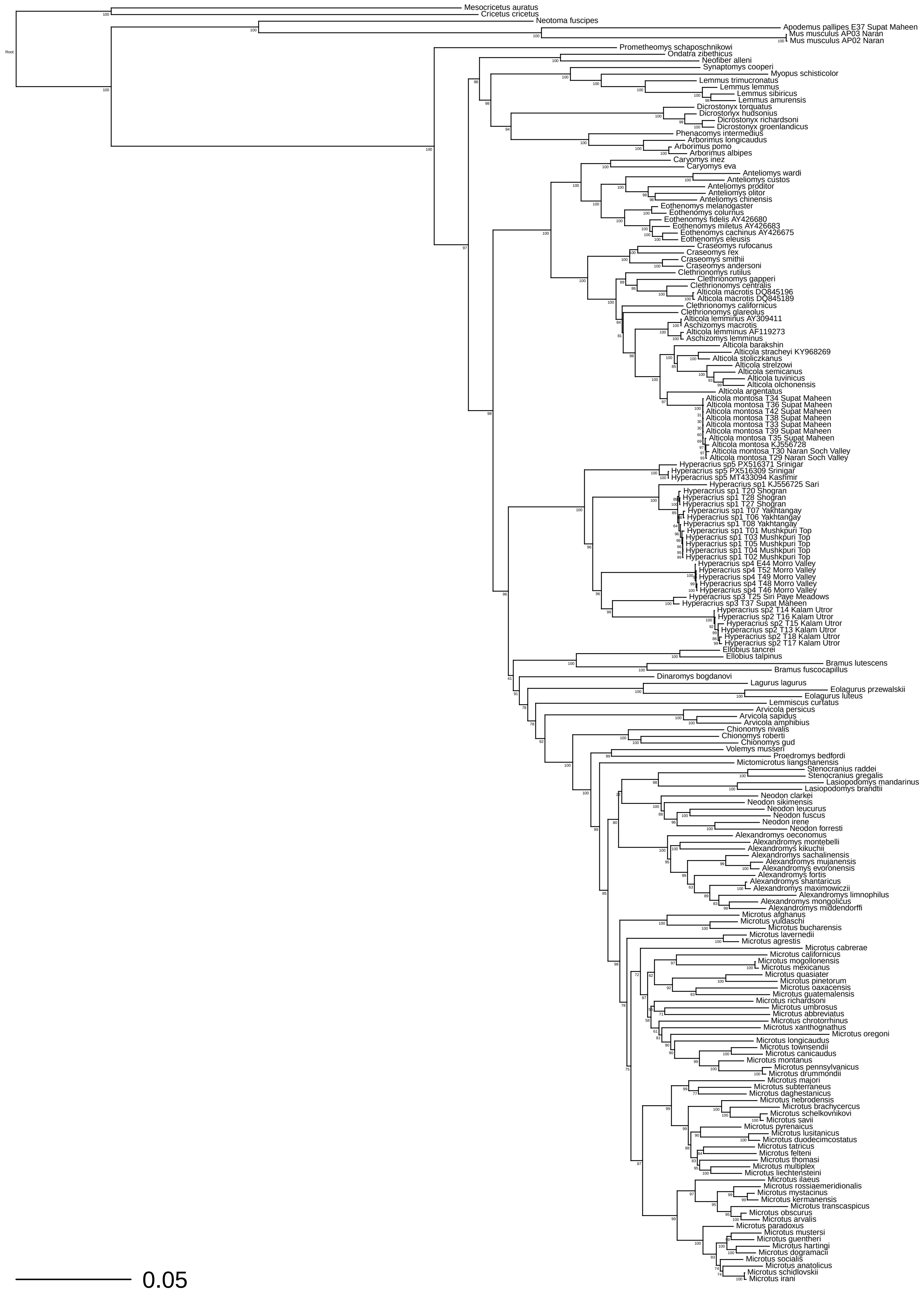
